# GIRK Channel Loss of Function Increases Dendritic Excitability in a Mouse Model of GNB1 Encephalopathy

**DOI:** 10.64898/2026.08.08.743706

**Authors:** Samuel Gritz, Anshul Voleti, Matthew S. Scarnati, Alessandro R. Galloni, Aaron D. Milstein

## Abstract

GNB1 encephalopathy (GNB1-E) is a rare neurodevelopmental disorder associated with motor dysfunction, epilepsy and learning disability caused by mutations in the gene encoding the G protein subunit Gβ_1_. Previous work has shown that altered Gβ_1_ can disrupt activation of G-protein-coupled inwardly rectifying potassium (GIRK) channels, dysregulate neuronal excitability and cause seizures. However, the relevant upstream regulators of Gβ_1_ and the consequences of GIRK dysfunction for neuronal synaptic, cellular and circuit function have not been characterized. Here we report that mice of both sexes carrying the deleterious p.I80T mutation in *Gnb1* present features consistent with GNB1-E, including developmental delay, decreased locomotion and increased anxiety. Using histology, whole-cell patch-clamp electrophysiology and pharmacology in *ex vivo* brain slices, we find that hippocampal neurons in heterozygous *Gnb1*^I80T/+^ mice exhibit simplified dendritic morphologies, decreased synaptic inhibition mediated by metabotropic GABA_B_ receptors and increased dendritic excitability. These phenotypes result in longer duration dendritic calcium spikes in response to synaptic afferent stimulation, an effect that is reversed by a specific activator of GIRK channels, ML297. Given the known roles of dendritic calcium spikes in driving burst firing and inducing synaptic plasticity, these findings suggest that targeting dendritic excitability has therapeutic potential to address both the seizure susceptibility and learning deficits associated with GNB1-E.

**Significance Statement:** GNB1 encephalopathy (GNB1-E) is a rare neurodevelopmental disorder associated with motor dysfunction, epilepsy and learning disability for which there are currently no mechanism-based treatments. Here we show that a pathogenic variant of the G protein subunit Gβ_1_ impairs activation of neuronal G-protein-coupled inwardly rectifying potassium (GIRK) channels by inhibitory synaptic GABA_B_ receptors. This leads to increased dendritic excitability and longer duration dendritic calcium spikes in mouse hippocampal neurons in response to stimulation of synaptic inputs. We find that this phenotype is reversed by a drug that activates GIRK channels, opening pathways to develop therapies for GNB1-E that specifically target dendritic excitability.

## Introduction

GNB1 encephalopathy (GNB1-E) is a rare autosomal dominant neurodevelopmental disorder typically caused by *de novo* mutations in the gene encoding G protein subunit Gβ_1_. Characteristic clinical features include delayed development; muscle hypotonia, hypertonia and/or dystonia; abnormal electroencephalogram (EEG) waveforms and seizures; abnormal brain structure; and various neurological symptoms, such as intellectual disability, anxiety and depression (Petrovski et al., 2016; Steinrucke et al., 2016; Lohmann et al., 2017; Hemati et al., 2018; Szczałuba et al., 2018; Endo et al., 2020; Schultz-Rogers et al., 2020; Da Silva et al., 2021; Lansdon and Saunders, 2021; Reyes et al., 2023; Tsuji et al., 2023; Nasvytis et al., 2024). A recent proteomic study identified changes in the expression of Gβ_1_ in the hippocampus and cortex as common across patients with diverse epilepsy disorders (Pires et al., 2021), suggesting that understanding the mechanisms disrupted in GNB1-E could inform treatments for a broader set of conditions.

While Gβ_1_ is a fundamental component of G-protein-coupled receptor (GPCR) signaling pathways that regulate numerous cellular processes throughout the brain and body (Ford et al., 1998; Okae and Iwakura, 2010; Qayum et al., 2026), previous work has shown that multiple known pathogenic *GNB1* variants specifically disrupt the interaction between Gβ_1_ and G-protein-coupled inwardly rectifying potassium (GIRK) channels (Signorini et al., 1997; Mark and Herlitze, 2000; Dascal and Kahanovitch, 2015) while leaving intact interactions with voltage-gated calcium channels and GPCRs like D_2_ dopamine receptors (Dascal, 2001; Oldham and Hamm, 2008; Betke et al., 2012; Zamponi and Currie, 2013; Reddy et al., 2021; Colombo et al., 2023). In particular, one of the more common pathogenic *GNB1* variants in humans, p.I80T (Petrovski et al., 2016; Hemati et al., 2018; Endo et al., 2020; Tsuji et al., 2023), has been shown to decrease activation of GIRK channels in heterologous cells (Reddy et al., 2021) and to cause epilepsy in mice (Reddy et al., 2026), though its effects on cellular and synaptic function have not been investigated in mammalian neurons. In this study, we first verify that heterozygous *Gnb1*^I80T/+^ mice exhibit characteristic features of GNB1-E, including developmental delay, locomotor deficits and anxiety, and then investigate its impacts on neuronal physiology.

In the brain, GIRK channels are essential for mediating slow synaptic inhibition downstream of G-protein-coupled GABA_B_ receptors (Dutar and Nicoll, 1988; Dutar et al., 2000; Luscher and Slesinger, 2010). In hippocampal neurons, colocalization of GIRK channels and GABA_B_ receptors is highest in dendrites (Koyrakh et al., 2005; Degro et al., 2015), and GIRK channels have been shown to modulate nonlinear integration of excitatory and inhibitory synaptic inputs in dendrites (Malik and Johnston, 2017), suggesting that this fundamental neuronal computation may be disrupted by dysregulation of GIRK channel function in GNB1-E. Furthermore, the balance between dendritic excitation and inhibition is known to regulate dendritic calcium spikes, which are involved in the induction of synaptic plasticity and goal-directed learning (Takahashi and Magee, 2009; Lovett-Barron et al., 2012; Bittner et al., 2015; Milstein et al., 2015; Bittner et al., 2017; Grienberger et al., 2017; Magee and Grienberger, 2020; Grienberger and Magee, 2022; Rolotti et al., 2022; Li et al., 2024; Madar et al., 2025; Udakis et al., 2025; Xiao et al., 2025; Yaeger et al., 2025; Campbell et al., 2026; Magee, 2026; Vaasjo et al., 2026). Accordingly, disruption of dendritic structure and function has been implicated in underlying cognitive deficits associated with epilepsy and neurodevelopmental disorders (Brager and Johnston, 2014; Copf, 2016; Prem et al., 2020; Nelson and Bender, 2021; Masala et al., 2023; Prem et al., 2024). Therefore, in this study we focused on the impact of *Gnb1* mutation in mice on synaptic transmission and dendritic excitability. We found that in heterozygous *Gnb1*^I80T/+^ mice, hippocampal pyramidal neurons exhibit fewer dendritic branches, reduced synaptic inhibition mediated by GABA_B_ receptor activation of GIRK channels, and increased propensity and duration of dendritic calcium spikes. The dendritic hyperexcitability phenotype was mimicked in wild-type (WT) neurons by pharmacological blockade of GIRK channels with ethosuximide (Kobayashi et al., 2009) and reversed in *Gnb1*^I80T/+^ neurons by the GIRK activator ML297 (Kaufmann et al., 2013; Wydeven et al., 2014; Huang et al., 2018). These findings suggest that targeting the mechanisms that regulate dendritic calcium spikes may offer therapeutic benefits for patients with GNB1-E.

## Materials and Methods

### Mice

All animal procedures and experiments were approved by the Rutgers University Institutional Animal Care and Use Committee (IACUC; protocol # 202200043). All animals were housed within facilities that are accredited by the Association for the Assessment and Accreditation of Laboratory Animal Care (AAALAC) and under the care of a veterinarian, in compliance with all relevant ethical regulations for the use of animals. Animals were housed in large cages with a 12:12 hour light-dark cycle and fed ad libitum. *Gnb1*^I80T/+^ mice were rederived from frozen sperm (JAX stock # 412228) using C57BL/6NJ females as hosts (Reddy et al., 2026). The genetic background strain of the source sperm and the resulting inbred offspring was a hybrid of C57BL/6NJ (JAX stock # 005304) and FVB.129P2 (JAX stock # 004828). *Gnb1*^I80T/+^ mice survived at slightly less than Mendelian ratios (Supp. Fig. S1). Between P8–P10, toes were clipped for identification, and the tissue was processed for genotyping by Transnetyx. Mouse pups were weaned and separated by sex and genotype at age P28. Age-matched WT littermates were used as controls for all experiments. Body weights were measured during genotyping (P8–P10), at weaning (P28) and at adulthood (∼7 weeks). To help offset delayed growth of *Gnb1*^I80T/+^ mice (Fig. 1A,B), the nutrition of all mice was supplemented with a high-calorie gel diet (ClearH20 DietGel Boost). To reduce anxiety in male mice, some partially soiled bedding was left unreplaced during bedding changes.

**Figure 1.**
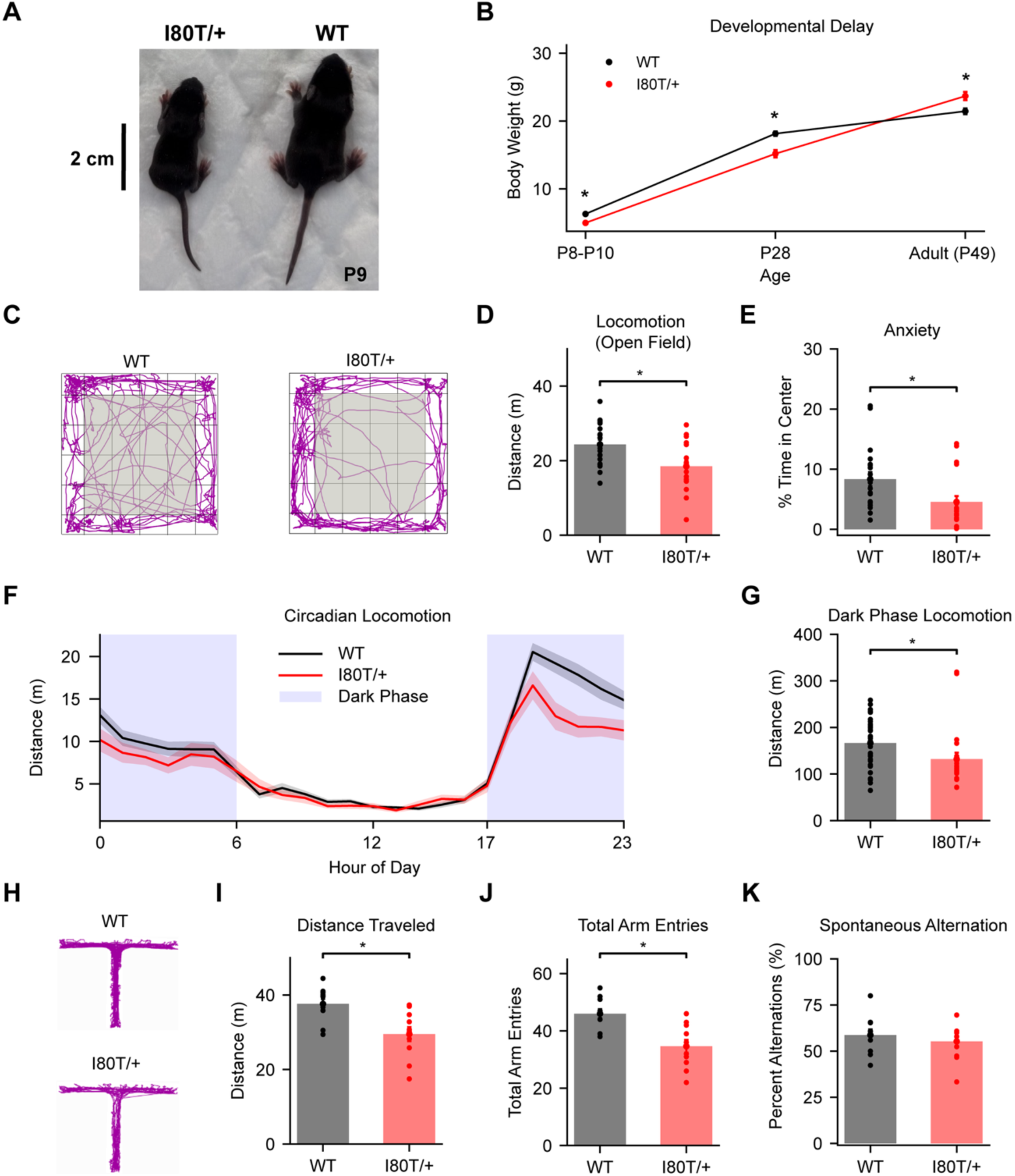
***A***, Representative photo of a female *Gnb1*^I80T/+^ mouse (left) and a female WT littermate (right) at P8. ***B***, Body weight during development (WT: black, *Gnb1*^I80T/+^: red). P8–P10: WT (*n* = 41): 6.30 ± 0.15 g vs. *Gnb1*^I80T/+^ (*n* = 36): 4.98 ± 0.13 g (*U* = 1256, *p* < 0.0001); P28: WT (*n* = 42): 18.14 ± 0.32 g vs. *Gnb1*^I80T/+^ (*n* = 38): 15.18 ± 0.55 g (*U* = 1330, *p* < 0.0001); Adult: WT (*n* = 32): 21.44 ± 0.42 g vs. *Gnb1*^I80T/+^ (*n* = 31): 23.68 ± 0.61 g (*U* = 293, *p* = 0.0054). ***C***, Representative locomotion traces during open field exploration (WT: left, *Gnb1^I80T/+^*: right). Grey shaded area indicates center zone. ***D***, Distance traveled during open field exploration. WT (*n* = 20): 24.36 ± 1.26 m vs. *Gnb1*^I80T/+^ (*n* = 20): 18.49 ± 1.37 m (*U* = 304, *p* = 0.0051). ***E***, Anxiety during open field exploration quantified as the percentage of time spent in the center zone. WT (*n* = 20): 8.33 ± 1.16% vs. *Gnb1*^I80T/+^ (*n* = 20): 4.56 ± 0.96% (*U* = 304, *p* = 0.0051). ***F***, Distance traveled per hour during diurnal locomotion in home cage, averaged across days. Shading reflects SEM across animals. Blue background marks circadian dark phases (WT: *n* = 32, *Gnb1*^I80T/+^: *n* = 22). ***G***, Distance traveled per day during waking (dark phase) hours (same data as ***F***). WT: 166.47 ± 9.20 m vs. *Gnb1*^I80T/+^: 132.07 ± 13.91 m (*U* = 514, *p* = 0.0045). ***H***, Representative locomotion traces during T-maze exploration. ***I***, Distance traveled during T-maze exploration. WT (*n* = 11): 37.61 ± 1.34 m vs. *Gnb1*^I80T/+^ (*n* = 11): 29.51 ± 1.88 m (*U* = 106, *p* = 0.0031). ***J***, Total number of left and right port entries during T-maze exploration. WT (*n* = 11): 46.00 ± 1.56 vs. *Gnb1*^I80T/+^ (*n* = 11): 34.64 ± 2.26 (*U* = 97, *p* = 0.0180). ***K***, Percentage of spontaneous left-right port alternations during T-maze exploration. WT (*n* = 11): 58.72 ± 3.04% vs. *Gnb1*^I80T/+^ (*n* = 11): 55.28 ± 2.95% (*U* = 70.5, *p* = 0.5315). All data represented as mean ± SEM. Statistics reflect Mann-Whitney U-tests. Asterisks indicate *p* < 0.05.

### Mouse behavior

#### Open field locomotion and anxiety

Open field exploration tests were performed on 7–12-week-old single-housed male and female mice. Animals were handled for 1 minute daily over the course of 3–5 days by the same experimenter prior to testing, and a handling tunnel was used to transfer mice into the arena to minimize stress (Gouveia and Hurst, 2017). Cages were transferred to the testing room to acclimate for at least 30 minutes before testing. Testing was performed in dim light in a covered plexiglass arena (50 cm × 50 cm × 40 cm). Each mouse was placed facing the wall in the same release corner and allowed to explore freely for 5 minutes. Time spent in the outer and center zones, and total distance traveled were measured using ANY-maze software (Stoelting). An anxiety metric was calculated as the percentage of total exploration time spent in the center zone (Crawley, 1985). The arena was cleaned with 70% ethanol after each mouse.

#### Circadian locomotor activity

Immediately following weaning (P28), single-housed male and female mice were transferred to a Digital Ventilated Cage (DVC^®^) system (Tecniplast) that provided 24-hour monitoring and tracking of animal well-being and locomotor activity via 12 capacitance-sensing electrodes positioned on the cage’s floor (Collins et al., 2025; Tir et al., 2025). The locomotor activity of mice was recorded from 4–7 weeks old while they were subject to a 12:12 hour light-dark cycle. Hourly tracking distance data was downloaded from DVC® Analytics software and processed with custom Python code for analysis and visualization (Gritz and Milstein, 2026).

#### T-maze exploration

To evaluate spatial working memory, spontaneous alternation tests in a T-maze were performed on 7–12-week-old single-housed male and female mice (d’Isa et al., 2021). A custom designed T-maze with removable doors and high walls (San Diego Instruments) was used to minimize the ability of anxious mice to exit the apparatus during testing. A handling tunnel was used to transfer mice into the maze start zone to minimize stress (Gouveia and Hurst, 2017). Mice were allowed to explore freely for 10 minutes, and their location was tracked using ANY-maze software (Stoelting). Spontaneous alternation behavior was analyzed using custom Python code (Gritz and Milstein, 2026). The animal’s path was recorded as a discrete sequence of zone entries (Start, Center, Left Arm and Right Arm). The position sequence was segmented into individual trips, defined as Start-to-Start intervals, and the first arm visited (Left or Right) during each trip was identified. An alternation was defined as a consecutive pair of trips in which the animal chose a different arm than on the immediately preceding trip. The percentage of spontaneous alternations was calculated as the number of alternating consecutive trip-pairs divided by the total number of consecutive trip-pairs (total arm choices minus one), multiplied by 100. The T-maze was cleaned with 70% ethanol after each mouse.

#### Hippocampal slice preparation

6–12-week-old male and female mice were deeply anesthetized with isoflurane and perfused transcardially with an ice-cold cutting solution containing: 210 mM sucrose, 25 mM NaHCO_3_, 2.5 mM KCl, 1.25 mM NaH_2_PO_4_, 0.75 mM CaCl_2_, 7 mM MgCl_2_, 7 mM glucose, 3 mM Na pyruvate and 1 mM ascorbic acid; osmolarity 300–310 mOsm; pH 7.3–7.4. Brains were then rapidly extracted and sliced on a vibratome (Leica) while submerged in an ice-cold bath containing oxygenated cutting solution. To prepare longitudinal slices of hippocampal area CA1, dorsal-ventral blocking cuts were made in each brain hemisphere at a 45° angle to the midline, parallel to the long axis of the hippocampus, as previously described (Milstein et al., 2015). This procedure yields 1–2 slices (250 μm thick) per hemisphere that are composed entirely of area CA1 from the dorsal/anterior two-thirds of the hippocampus and are in plane with the somatodendritic axis of pyramidal neurons. Slices were immediately transferred to a submerged holding chamber containing heated (35°C) oxygenated artificial cerebrospinal fluid (ACSF): 125 mM NaCl, 25 NaHCO_3_, 2.5 mM KCl, 1.25 mM NaH_2_PO_4_, 1 mM MgCl_2_, 2.0 mM CaCl_2_, 25 mM glucose, 3 mM Na pyruvate and 1 mM ascorbic acid; osmolarity 300–310 mOsm; pH 7.3–7.4. The holding chamber was maintained at 35°C for 30 minutes, then kept at room temperature (RT; 22°C) for up to 4 hours. All solutions contained fresh pyruvate and ascorbic acid and were continuously bubbled with 95% O_2_ and 5% CO_2_. Additionally, prior to recording, the ventral/posterior third of each slice was excised with a scalpel.

#### Intracellular electrophysiology

During recording, slices were perfused continuously with heated, oxygenated ACSF (described above). In a subset of experiments, the following drugs were added to the perfusate: 10 μM gabazine (SR 95531 hydrobromide, Tocris, Bristol, UK; 10 mM stock dissolved in ultrapure water) to block GABA_A_ receptors, 10 μM Baclofen (Sigma; 10 mM stock dissolved in 10 mM HCl and ultrapure water) to activate GABA_B_ receptors, 10 μM ML297 (Sigma; 10 mM stock dissolved in DMSO) to activate GIRK channels, or 500 μM ethosuximide (Sigma; 10 mM stock dissolved in ultrapure water) to block GIRK channels. A Peltier heating element maintained the bath temperature at 35±1 °C. Pyramidal neurons in hippocampal area CA1 were visualized for intracellular patch-clamp recordings using a Dodt gradient contrast microscope (Scientifica) and a water-immersion objective (Olympus, 20x/0.5 NA or 60x/1.0 NA). Patch electrodes were pulled from borosilicate glass capillaries (GC150F-7.5, with filament, Harvard Apparatus) using a Zeitz DMZ Universal Electrode Pipette Puller to yield an electrical resistance of 6–8 MΩ. Recording electrodes were filled with an intracellular solution containing: 134 mM K gluconate, 6 mM KCl, 4 mM NaCl, 0.3 mM Tris⋅GTP, 4 mM Mg_2_ATP, 14 mM (di)Tris phosphocreatine, 10 mM HEPES, 0.5% w/v biocytin hydrochloride and 50 µM Alexa Fluor 488 hydrazide; osmolarity 290–295 mOsm; pH 7.3. Analog intracellular voltage signals were received by a Multiclamp 700B amplifier (Molecular Devices), dynamically filtered by a HumBug (Digitimer) to reduce 60 Hz electrical interference, and digitized using a National Instruments DAQ board. Voltage recordings were also monitored live using a BK Precision 2563-MSO Oscilloscope. During whole-cell current clamp recordings, (series) resistance was measured and corrected for through the Multiclamp bridge balance compensation circuit. Liquid junction potentials were not compensated. Recordings were rejected if neurons had a resting membrane potential >-50 mV, access resistance >40 MΩ, or if access resistance changed by more than 30% over the course of the recording. Access resistance was comparable across recordings of neurons from both genotypes (WT: *n* = 44, 31.14 ± 1.77 MΩ, *Gnb1*^I80T/+^: *n* = 44, 27.97 ± 1.49 MΩ, *U* = 1133, *p* = 0.1697, Mann-Whitney U-test). Input and access resistances were continuously measured using 100 ms, −50 pA current pulses. Electrical stimulation of afferent axons was performed using 0.1 MΩ impedance blunt-tipped bipolar platinum-iridium electrodes (75 μm spacing, MicroProbes, Gaithersburg, MD), fed through a glass capillary for stable positioning by a micromanipulator (Scientifica). Stimulating electrodes were placed in the stratum lacunosum-moleculare (SLM) and either the stratum radiatum (SR) or stratum oriens (SO) layers of hippocampal area CA1 ∼300-400 μm from the recorded cell to stimulate the perforant path (PP) axons from entorhinal cortex layer III (ECIII) and the Schaffer collateral (SC) inputs from hippocampal area CA3 onto the proximal apical or basal CA1 dendrites, respectively. Electrical stimulation pulses were generated by a multi-channel stimulus generator (STG5, Multi Channel Systems) using the MC Stimulus III software. Individual stimuli were biphasic, consisting of a 100 μs pulse followed by a 200 μs charge-balancing pulse with opposite polarity and half amplitude. Current pulse intensities (30-1400 μA) were calibrated for each individual stimulus pathway such that the average initial amplitude of recorded single-pulse excitatory postsynaptic potentials (EPSPs) was ∼2 mV before application of gabazine.

#### Electrophysiology data acquisition and analysis

Intracellular current injection waveform control and recording of intracellular voltage signals was performed using the Neuromatic plugin for the Igor Pro software (WaveMetrics) (Rothman and Silver, 2018). Further analysis and visualization of data was performed using custom Python code (Gritz and Milstein, 2026). Input resistance was calculated from the measured peak voltage response to brief (100 ms) hyperpolarizing current steps (−50 pA) using Ohm’s law. To assay hyperpolarization-activated cation current *I*_h_ (Maccaferri et al., 1993), voltage sag was quantified from voltage responses to larger (−200 pA) and longer duration (500 ms) hyperpolarizing current steps using the 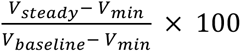, where *_Vmin_* is the peak hyperpolarization, *_Vsteady_* is the mean voltage in the final 10 ms of the current step, and *_Vbaseline_* is the mean voltage in the 10 ms prior to the current step onset. Resting membrane potential was recorded in the absence of current injection, and then resting potentials were monitored and maintained at −70 mV throughout the recording with an offset current injection. Action potential (AP) and after-hyperpolarization (AHP) properties were analyzed from the first spike elicited at rheobase. The AP threshold was defined as the voltage corresponding to the maximum of the second derivative of the membrane potential (*_d_*^2^*_V_*/*_dt_*^2^), or the first derivative if a second-derivative peak was not detected. AP size was measured as the difference between the peak and threshold voltages, and the AP halfwidth was defined as the duration at the voltage midpoint between threshold and peak. AHP amplitude was measured relative to AP threshold. AHP area was calculated as the integral of the threshold-subtracted voltage trace until recovery to zero, or within a maximum window duration of 200 ms after spike onset. F-I curve midpoints were estimated from a sigmoidal fit (Fellous et al., 2003). Spike rate adaptation was measured by first calibrating the current injection amplitude in each neuron to produce exactly six action potentials.

To quantify stimulus-frequency-dependent synaptic integration and excitatory-inhibitory (E/I) balance, we measured the peak amplitude of synaptic responses to trains of 3 stimuli delivered with the following inter-stimulus intervals (ISIs): 300, 100, 50, 25, 10 ms, before and after block of GABA_A_ receptors with gabazine (Milstein et al., 2015). This procedure was performed for both proximal (SC) and distal (PP) stimulating electrodes. Fast GABA_A_ receptor-mediated inhibitory postsynaptic potential (IPSP) amplitude was estimated from the difference between average compound EPSP waveforms before and after application of gabazine. E/I imbalance was quantified from the formula 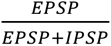. Following application of gabazine, GABA_B_ receptor-mediated IPSP amplitude and area was measured from the remaining slow hyperpolarizing component of the synaptic response within a window duration of 300 ms after stimulus onset. To quantify nonlinear summation, we first measured the unitary EPSP waveform by averaging across the individual stimuli delivered at 300 ms intervals in gabazine, followed by rectification of any hyperpolarization below baseline. Then, we computed the expected compound EPSP waveforms that would result from linear summation of the unitary EPSP at each of the stimulation frequencies tested. Supralinear summation was measured from the difference between expected and actual recorded compound EPSP waveforms.

To synaptically evoke dendritic calcium spikes (plateau potentials), proximal SC and distal PP pathways were stimulated either individually or simultaneously in a theta burst pattern consisting of 5 bursts delivered with an inter-burst interval of 150 ms, with each burst consisting of 5 pulses delivered with an ISI of 10 ms. This was performed in the presence of gabazine. To quantify plateau potentials, first fast action potentials were clipped, and the voltage waveform was linearly interpolated to estimate the underlying waveform of depolarization. Then, the plateau area was measured as the integral of the baseline-subtracted depolarization waveform wherever it exceeded a threshold of 20 mV (Bittner et al., 2015; Milstein et al., 2015).

#### Immunohistochemistry and analysis of neuronal morphology

After recording, slices were immediately fixed overnight at 4°C in a 4% formaldehyde solution and then stored in PBS. For immunohistochemical detection of neurons filled with biocytin, fixed slices were first incubated for 1–2 hr at RT in a blocking solution containing 0.5% Triton X-100 and 5% Normal Goat Serum (NGS) in PBS. Slices were washed twice (10 min each) in PBS and incubated overnight in a staining solution containing 0.05% Triton X-100, 0.5% NGS and 594-conjugated streptavidin (2 µg/ml) or 488-conjugated streptavidin (1 µg/ml). Slices were then washed twice (10 minutes each) in PBS and then mounted on glass slides. Images were acquired using a confocal microscope (Zeiss LSM900 Laser Scanning Confocal Microscope, objective: 20x/0.7NA, pinhole size: 1 airy unit). The images were used to reconstruct the apical and basal dendrites of recorded CA1 pyramidal neurons using the Simple Neurite Tracer (SNT) plugin in Fiji (Schindelin et al., 2012; Arshadi et al., 2021). Following full reconstruction, quantification of the number and spatial distribution of dendritic branches (Sholl analysis) (Sholl, 1953) was performed using the SNT Fiji plugin and custom Python code (Gritz and Milstein, 2026).

#### Biochemistry

Acute P7 mouse hippocampal brain slices were rapidly collected and snap-frozen on dry ice. Western blotting was performed to measure Gβ_1_ protein in *Gnb1*^I80T/+^ and WT mice. Cells were lysed in Pierce RIPA lysis buffer (ThermoFisher) with HALT protease and phosphatase inhibitor (ThermoFisher). Slices were resuspended in 500 µL of ice-cold lysis buffer and transferred to a 1.5 mL microcentrifuge tube containing 2.8 mm stainless steel beads. Tissue was homogenized using a BeadBug microtube homogenizer (Benchmark Scientific) at high speed for 2 cycles of 30 seconds each, with cooling on ice between cycles to prevent overheating. Homogenates were incubated at 4 °C with constant rotation for 1 hour, followed by centrifugation at 14,000 x g for 20 minutes at 4 °C to pellet debris. The clarified supernatant was collected, and protein concentration was determined using a BCA assay. Lysates were then used for SDS-PAGE and western blot analysis. Protein lysates (50 µg) were separated on a 4-12% acrylamide SDS-PAGE gel, followed by a 1-hour transfer to nitrocellulose membranes. Membranes were blocked in Intercept (TBS) blocking buffer (LI-COR Biosciences) for 1-hour and incubated in rabbit anti-Gβ_1_ (1:1000, Cell Signaling Technology) and mouse anti-Vinculin (Cell Signaling Technology) antibodies overnight at 4 °C. The following day, membranes were washed 3x (5 min each) in TBS with Tween20 (TBST), incubated in IRDYE 680RD Goat anti-

Rabbit IgG secondary antibody and in IRDYE 800CW Goat anti-Mouse IgG secondary antibody (1:10,000; LI-COR Biosciences) for 1-hour at RT. Blots were then washed 3 times (5 min each) in TBST. Blots were imaged using the Odyssey M Imaging System (LI-COR Biosciences) according to the manufacturer’s instructions. Band intensities were quantified using Image Studio software (LI-COR Biosciences), and Gβ_1_ protein level was normalized to Vinculin prior to statistical analysis. Gβ_1_ protein levels in *Gnb1*^I80T/+^ mice were indistinguishable from WT (Supp. Fig. S2), suggesting that the observed phenotypes are due to altered protein function rather than haploinsufficiency.

#### Statistical analysis

All statistical analysis was performed using custom code (Gritz and Milstein, 2026) using the scipy.stats package (Virtanen et al., 2020) in Python and the lme4 (Bates et al., 2015), lmerTest (Kuznetsova et al., 2017), emmeans (Lenth and Piaskowski, 2026), car (Fox and Weisberg, 2019) and nlme (Pinheiro et al., 2026) packages in R (Giorgi et al., 2022). Simple comparisons between two groups of samples (e.g., behavioral metrics, cellular input resistance) were performed using nonparametric tests: the Mann-Whitney U test for unpaired data, the Wilcoxon Signed-Rank test for paired data, and the Kolmogorov-Smirnov test for cumulative distributions. All statistical tests were two-sided, and p-values < 0.05 were considered statistically significant. To account for complex experimental designs with repeated measures or hierarchical data structures (e.g., E/I balance, supralinear summation), we employed linear mixed effects (LME) modeling. Models were specified with the following structure: Response ∼ Genotype + Genotype × ISI + (1 | Subject), where Genotype and its interaction with ISI are fixed effects, and Subject is a unique ID per animal or per cell that was included as a random effect to account for within-subject correlations. This model tests whether there is a main effect of Genotype (offset), and whether there is an interaction between Genotype and ISI, meaning the slope of the relationship between Response and ISI differs between Genotypes. This model structure was used to compare responses across genotypes given the same stimulus pathway and drug condition. Significance of fixed effects and interactions was determined using Type III Analysis of Variance with Satterthwaite’s method for degrees of freedom approximation. When an interaction effect (e.g., Genotype × ISI) was statistically significant (p < 0.05), post-hoc pairwise comparisons were performed to identify which specific ISIs showed significant differences between genotypes. These post-hoc comparisons were performed as pairwise t-tests using estimated marginal means. The resulting p-values were then corrected for multiple comparisons using the Benjamini-Hochberg False Discovery Rate (FDR) method (Benjamini and Hochberg, 1995). When an interaction effect was significant, but individual post-hoc comparisons did not reach a significance threshold of *_α_* < 0.05 after FDR correction, this indicates an interaction effect between genotype and ISI was distributed across all ISIs tested rather than concentrated at a subset of ISIs. In the figures, significant main effects were indicated by asterisks bracketing all inter-stimulus interval conditions, significant interaction effects were indicated by hashtag symbols (#) bracketing all inter-stimulus interval conditions, and significant post-hoc comparisons at individual ISIs were marked with asterisks. For f-I curve and spike frequency adaptation analyses, 2-way Repeated Measures ANOVA tests were used. Specific statistical tests and sample sizes for each experiment are reported in the figure legends.

## Results

### *Gnb1*^I80T/+^ mice exhibit behavioral phenotypes consistent with GNB1 encephalopathy

Patients with GNB1-E, including those with the p.I80T variant of *GNB1*, present with motor dysfunction, developmental delay, and behavioral abnormalities (Petrovski et al., 2016; Hemati et al., 2018). Thus, before using *Gnb1*^I80T/+^ mice as a model system to investigate the synaptic and cellular mechanisms of the disorder, we first sought to determine whether *Gnb1*^I80T/+^ mice recapitulate these phenotypes. We first measured their body weights over the timecourse of development and observed that *Gnb1*^I80T/+^ pups had significantly lower body size and weight than their WT littermates at P8–P10 and P28 (Fig. 1A,B), consistent with delayed development. By adulthood *Gnb1*^I80T/+^ mice had caught up to and slightly exceeded the body weights of control mice.

Next, we assessed spontaneous exploratory locomotor activity in an open field arena. Adult *Gnb1*^I80T/+^ mice were significantly less active than WT littermates, covering a shorter total distance during 5 minutes of exploration (Fig. 1C,D). To assess anxiety-like behavior, we measured the time spent in the center versus the outer zone of the open field. Adult *Gnb1*^I80T/+^ mice spent less time than control mice in the center of the arena (Fig. 1C,E), consistent with heightened anxiety (Crawley, 1985). We also analyzed home cage circadian locomotor activity and found that 4–7-week-old *Gnb1*^I80T/+^ mice exhibited reduced locomotion during their most active (dark) hours (Fig. 1F,G). Finally, we assessed spatial working memory by measuring spontaneous exploration behavior in a T-maze. Consistent with the reduced locomotor activity in the open-field arena and the home cage (Fig. 1C–G), 7–12-week-old *Gnb1*^I80T/+^ mice covered a shorter total distance in the T-maze (Fig. 1H,I), and entered the left and right arms of the T maze fewer times than WT mice (Fig. 1J). During T-maze exploration, alternating consecutive choices to turn left or right from the center arm is a measure of short-term spatial working memory (d’Isa et al., 2021). According to this metric, *Gnb1*^I80T/+^ mice did not exhibit a deficit in spatial working memory (Fig. 1K). In summary, *Gnb1*^I80T/+^ mice exhibit features consistent with GNB1-E, including delayed body development, decreased motor function and anxiety-like behavior, supporting its use as a model system to investigate the impact of *Gnb1* mutation on cellular and synaptic function in mammalian neurons.

### Decreased somatic excitability in *Gnb1^I80T/+^* neurons

To assess changes in neuronal excitability resulting from *Gnb1* mutation, we performed whole-cell patch-clamp recordings from pyramidal neurons in acute slices of hippocampal area CA1 from 6–12-week-old male and female *Gnb1*^I80T/+^ and WT mice. A number of physiological parameters tested were not different between genotypes, including resting membrane potential, the voltage threshold for action potentials, the size and shape of action potentials (APs), and the area of spike after-hyperpolarizations (AHPs) (Table 1; Fig. 2). We also assayed hyperpolarization-activated cation current *I*_h_ from the dynamic response to hyperpolarizing current injection (voltage sag) (Maccaferri et al., 1993) and measured spike-rate adaptation during brief step current injections, and found no differences between genotypes (Fig. 2B, H, I). However, several parameters measured were consistent with an unexpected decrease in excitability in *Gnb1*^I80T/+^ neurons, including decreased input resistance, increased current threshold for action potentials (rheobase), and a rightward shift in the relationship between injected current amplitude and output firing rate (f-I curve) (Table 1; Fig. 2A, C–G). This observed decrease in neuronal excitability is inconsistent with increased seizure susceptibility, suggesting that if *Gnb1*^I80T/+^ neurons do have a loss of GIRK function, any resulting increase in excitability has been compensated for by up-regulating other leak or voltage-gated conductances, resulting in a net decrease in excitability measured at the soma.

**Figure 2.**
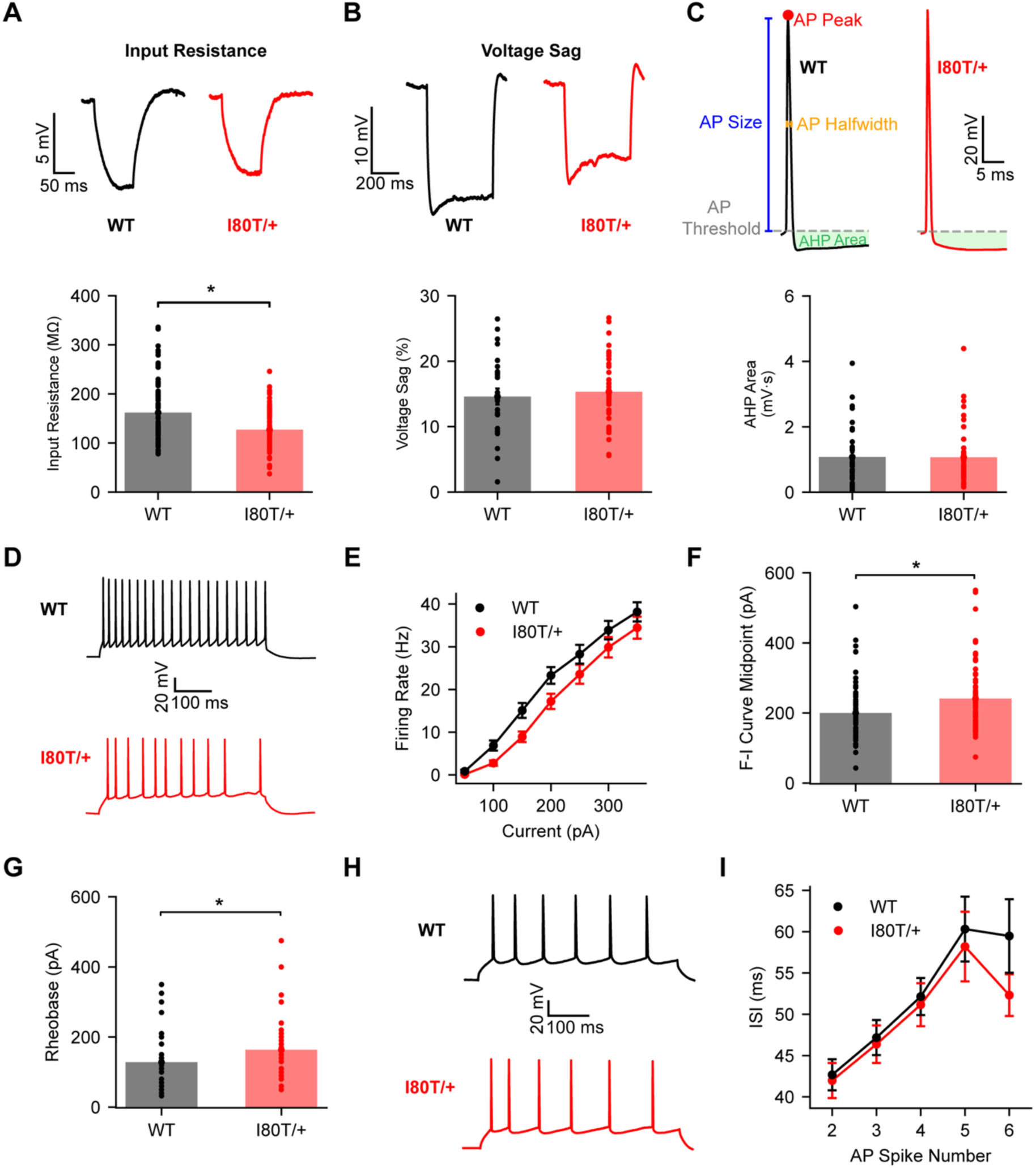
***A***, Representative voltage traces recorded during brief (100 ms) hyperpolarizing current injections (top) to measure input resistance (bottom). WT (*n* = 90): 161.84 ± 6.41 MΩ vs. *Gnb1*^I80T/+^ (*n* = 88): 127.03 ± 4.32 MΩ (*U* = 5296.5, *p* = 0.0001). ***B***, Representative voltage traces recorded during extended (500 ms) hyperpolarizing current injections (top) to measure voltage sag (bottom). WT (*n* = 26): 14.60 ± 1.21 vs. *Gnb1*^I80T/+^ (*n* = 44): 15.32 ± 0.74 (*U* = 521.5, *p* = 0.5434). ***C***, Top: representative action potential and after-hyperpolarization (AHP) waveforms recorded in response to current injections at rheobase. Bottom: AHP area. WT (*n* = 37): 1.08 ± 0.14 mV * s vs. *Gnb1*^I80T/+^ (*n* = 45): 1.07 ± 0.13 mV * s (*U* = 832, *p* = 1.000). ***D***, Representative voltage traces recorded during suprathreshold depolarizing current injections. ***E***, Spike frequency-current relationship (f-I curve). WT (*n* = 68) vs. *Gnb1*^I80T/+^ (*n* = 70); 2-way repeated-measures ANOVA; main effect of genotype: *F*(1,123) = 2.649, *p* = 0.1062; genotype × current interaction effect: *F*(14,1034) = 1.16, *p* = 0.3008. ***F***, The mid-point of each f-I curve from ***E*** was calculated from a sigmoidal fit. WT (*n* = 68): 200.20 ± 10.0 pA vs. *Gnb1*^I80T/+^ (*n* = 70): 241.10 ± 11.30 pA (*U* = 1682.5, *p* = 0.0030). ***G***, Rheobase current amplitude. WT (*n* = 37): 128.30 ± 13.10 pA vs. *Gnb1*^I80T/+^ (*n* = 45): 163.40 ± 12.70 pA (*U* = 592, *p* = 0.0245). ***H***, Representative voltage traces recorded during depolarizing current injections calibrated to produce exactly six spikes to analyze spike rate adaptation. ***I***, Adaptation of inter-spike interval (ISI) with increasing spikes in a train. 2-way repeated-measures ANOVA; main effect of genotype: F(1,118) = 0.1157, *p* = 0.7344); genotype × spike number interaction effect: F(4,472) = 0.0719, *p* = 0.9906. All data represented as mean ± SEM. Statistics reflect Mann-Whitney U-tests unless otherwise stated. Asterisks indicate *p* < 0.05.

**Table 1.** Neuronal excitability properties measured intracellularly.

| **Table 1.**<br>Neuronal excitability properties measured intracellularly. |  |  |  |  |
| --- | --- | --- | --- | --- |
| **Property** | **WT** | **I80T/+** | ***U***<br>**statistic** | ***p*-value** |
| Resting membrane potential (mV) | -66.28 ± 0.59 (104) | -66.32 ± 0.64 (91) | 4786 | 0.8914 |
| Input resistance (MΩ) | 161.84 ± 6.41 (90) | 127.03 ± 4.32(88) | 5296.5 | **0.0001\*** |
| Voltage sag (%) | 14.60 ± 1.21 (26) | 15.32 ± 0.74 (44) | 521.5 | 0.5434 |
| Rheobase (pA) | 128.28 ± 13.15 (37) | 163.43 ± 12.69 (45) | 592 | **0.0245\*** |
| AP voltage threshold (mV) | -45.94 ± 0.97 (49) | -46.18 ± 0.96 (52) | 1221.5 | 0.7238 |
| AP size (mV) | 82.81 ± 1.97 (49) | 81.65 ± 2.01 (52) | 1342 | 0.6465 |
| AP halfwidth (ms) | 0.98 ± 0.03 (49) | 0.94 ± 0.03 (52) | 1336.5 | 0.6717 |
| AHP amplitude (mV) | 9.77 ± 0.64 (37) | 9.12 ± 0.89 (45) | 1002 | 0.1153 |
| AHP area (mV * s) | 1.08 ± 0.14 (37) | 1.07 ± 0.13 (45) | 832 | 1.000 |
| Data represented as mean ± SEM. Values in parentheses represent number of recorded cells. Statistics reflect Mann-Whitney U-tests. Asterisks and bold text indicate $p < 0.05$. | | | | |

### Altered dendritic morphology in *Gnb1*^I80T/+^ neurons

Changes in neurite outgrowth have recently been observed as a cellular phenotype associated with neurodevelopmental disorders (Copf, 2016; Prem et al., 2020; Prem et al., 2024). Thus, we investigated whether *Gnb1* mutation alters neuronal morphology. Each recorded neuron was filled with an intracellular marker (biocytin) to enable downstream histology, imaging, and morphological reconstruction and analysis. Example reconstructions are shown in Fig. 3A and Supp. Fig. S3. Sholl analysis was used to analyze the complexity of the basal and apical dendritic arbors of pyramidal neurons and the spatial distribution of dendritic branches along the somatodendritic axis (Sholl, 1953). This revealed a decrease in the number of basal branches, and a shift in the location of apical branches from distal to proximal in *Gnb1*^I80T/+^ neurons compared to WT (Fig. 3B– G). These layer-specific changes in dendritic complexity could translate to altered processing of specific synaptic inputs that differentially target basal and apical branches (Amaral and Witter, 1989; Spruston, 2008). Thus, we next investigated whether the synaptic integration function of dendrites is also impacted in *Gnb1*^I80T/+^ neurons.

**Figure 3.**
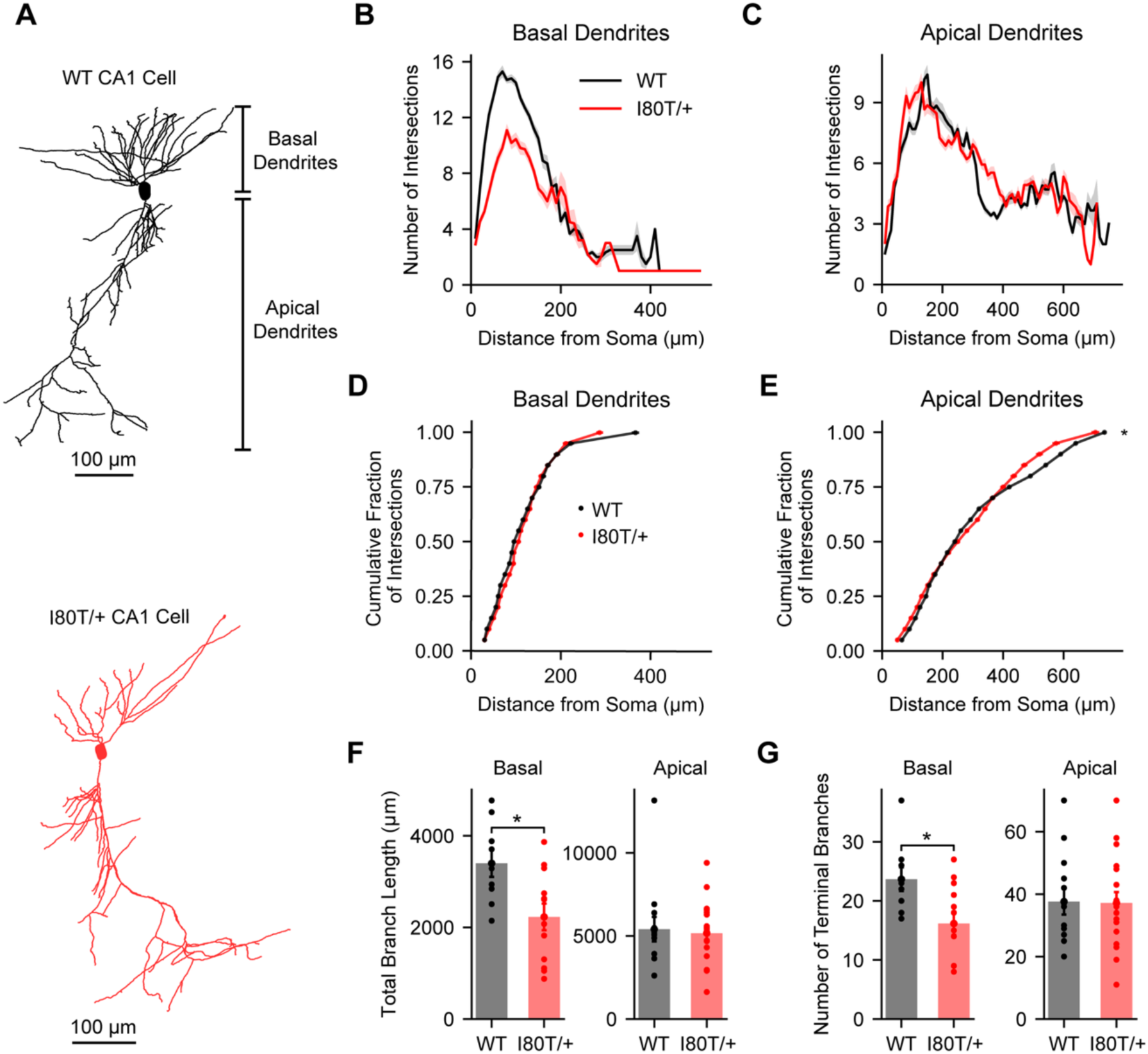
***A***, Representative neuronal morphologies histologically reconstructed following the intracellular recordings shown in Fig. 2. ***B***–***E***, Sholl analysis of dendritic branch distributions. ***B***, Number of dendritic branch intersections vs. distance from the soma for basal dendrites (WT: *n* = 9, *Gnb1*^I80T/+^: *n* = 12 neurons). Shading represents SEM across cells. ***C***, Same as ***B*** for apical dendrites (WT: *n* = 13, *Gnb1*^I80T/+^: *n* = 18 neurons). ***D***, Same data as ***B*** expressed as a cumulative fraction of dendritic branch intersections for basal dendrites. Kolmogorov-Smirnov test: *D =* 0.03546*, p* = 0.1480. ***E***, Same as ***D*** for apical dendrites (same data as ***C***). Kolmogorov-Smirnov test: *D* = 0.03758, *p* = 0.0023. ***F***, Total dendritic branch length (same neurons as ***B***–***E***). Left: basal branches. WT: 3399.94 ± 297.16 μm vs. *Gnb1*^I80T/+^ 2229.35 ± 291.07 μm (*U* = 86, *p* = 0.0252). Right: apical branches. WT: 5404.58 ± 736.94 μm vs. *Gnb1*^I80T/+^ 5167.66 ± 455.67 μm (*U* = 108, *p* = 0.7336). ***G***, Number of terminal branches. Left: basal branches. WT: 23.67 ± 2.00 vs. *Gnb1*^I80T/+^ 16.17 ± 2.0 (*U* = 83, *p* = 0.0422). Right: apical branches: WT: 37.62 ± 4.18 vs. *Gnb1*^I80T/+^ 37.17 ± 3.49 (*U* = 116, *p* = 0.9840). In ***F***–***G***, statistics reflect Mann-Whitney U-tests. All data represented as mean ± SEM. Asterisks indicate *p* < 0.05.

### Pathway-specific changes in excitatory and inhibitory synaptic integration in *Gnb1*^I80T/+^ neurons

In most mammalian hippocampal and cortical neural circuits, excitatory axonal projections from different brain regions and circuit layers form synapses onto specific subregions of the dendrites of pyramidal neurons (DeFelipe et al., 2002; Spruston, 2008; Petreanu et al., 2009; Larkum, 2013; Luo, 2021). These afferents also synapse onto various subclasses of inhibitory interneurons, which in turn synapse onto specific dendritic domains (Freund and Buzsaki, 1996; Gulyas et al., 1999; Megias et al., 2001; Palmer et al., 2012; Fino et al., 2013; Milstein et al., 2015; Bloss et al., 2016; Booker and Vida, 2018). In hippocampal area CA1, pyramidal neurons receive two primary excitatory afferents – inputs from hippocampal area CA3 to basal and proximal apical dendrites, and inputs from entorhinal cortex layer III (ECIII) to the distal apical dendrites (often referred to as “apical tuft” dendrites) (Amaral and Witter, 1989; Magee, 2000; Takahashi and Magee, 2009; Bittner et al., 2015). Here we investigated the relative strengths of mono-synaptic excitation and di-synaptic inhibition recruited by these two pathways onto three dendritic compartments (basal, proximal apical, and distal apical) using electrical stimulation of axons in *Gnb1*^I80T/+^ and WT slices (Milstein et al., 2015). We recorded intracellular postsynaptic potentials (PSPs) evoked by low frequency stimulation (300 ms inter-stimulus interval) both before and after application of gabazine, an antagonist of GABA_A_ receptors (Fig. 4A, see Methods). The strength of excitatory inputs was estimated from the peak amplitude of the average fast excitatory postsynaptic potential (EPSP) recorded after the application of gabazine (Fig. 4A,B). To estimate the fast component of inhibition mediated by GABA_A_ receptors, we measured the peak amplitude of the difference trace obtained by subtracting the average PSP recorded before gabazine from that recorded after gabazine (Fig. 4A,C). Finally, the slow component of inhibition mediated by GABA_B_ receptors was quantified from the area of the hyperpolarizing component of the average PSP waveform recorded in gabazine (Smirnov et al., 1999) (Fig. 4A,D). This procedure was repeated for stimulating electrodes placed in three sublayers of CA1 – the distal stratum lacunosum (SLM) layer to excite the ECIII inputs to distal apical dendrites, the proximal stratum radiatum (SR) layer to excite the CA3 inputs to proximal apical dendrites, and the stratum oriens (SO) layer to excite the CA3 inputs to basal dendrites. We observed pathway-specific deficits in the recruitment of synaptic inhibition onto *Gnb1*^I80T/+^ neurons. Fast GABA_A_ inhibition was specifically reduced in response to stimulation of the CA3 inputs to basal dendrites (Fig. 4A,C), which may be a consequence of reduced inhibitory synapse number given the observed decrease in basal dendritic branch number and length (Fig. 3). Slow GABA_B_ inhibition was specifically reduced in response to stimulation of the ECIII inputs to distal apical dendrites (Fig. 4A,D), consistent with the hypothesis that *Gnb1* mutation disrupts the recruitment of hyperpolarizing GIRK channels by synaptic GABA_B_ receptors (Dutar and Nicoll, 1988; Mark and Herlitze, 2000; Dascal and Kahanovitch, 2015; Reddy et al., 2021), which is expected to preferentially impact inputs to the distal dendrites where GABA_B_ receptors and GIRK channels are most highly co-expressed (Degro et al., 2015).

**Figure 4.**
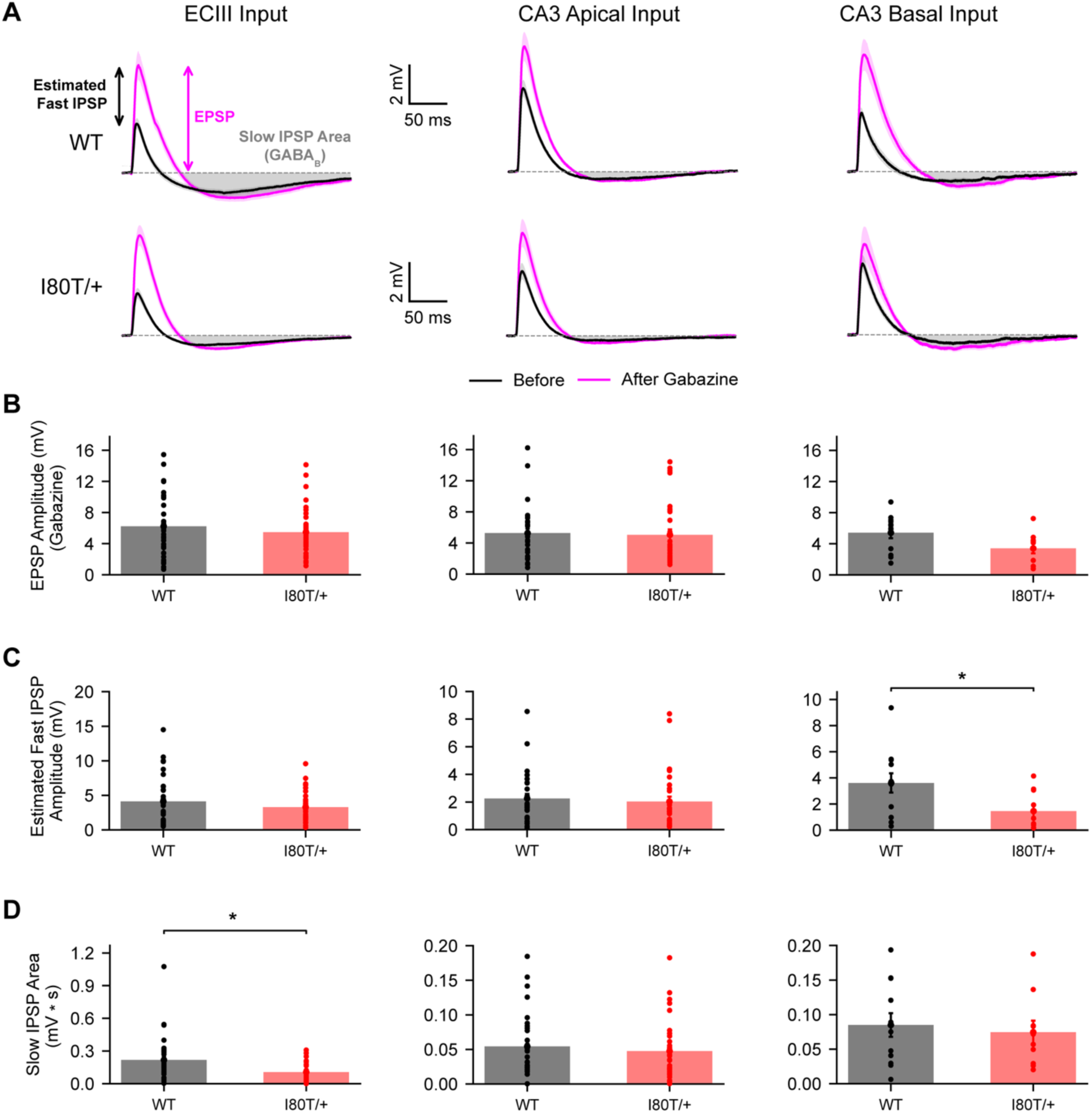
***A***, Traces show post-synaptic potentials (PSP) recorded before (black) and after application of gabazine (pink), allowing estimation of excitatory (EPSP), fast inhibitory (IPSP), and slow IPSP components (grey shaded area). Traces are averaged across 3 pulses stimulated with a 300 ms inter-stimulus interval (ISI), and then averaged across neurons. Shown are averaged traces from WT (top) and *Gnb1*^I80T/+^ (bottom) neurons for stimulation of each of three input pathways. Shading indicates SEM across neurons. ***B***, EPSP amplitude. Left: ECIII. WT (*n* = 33): 6.24 ± 0.71 mV vs. *Gnb1*^I80T/+^ (*n* = 34): 5.48 ± 0.54 mV (*U* = 602, *p* = 0.6115). Center: CA3 Apical. WT (*n* = 32): 5.26 ± 0.60 mV vs. *Gnb1*^I80T/+^ (*n* = 33): 5.05 ± 0.68 mV (*U* = 585, *p* = 0.4585). Right: CA3 Basal. WT (*n* = 12): 5.41 ± 0.70 mV vs. *Gnb1*^I80T/+^ (*n* = 10): 3.40 ± 0.63 mV (*U* = 88, *p* = 0.0698). ***C***, Estimated fast GABA_A_ IPSP amplitude. Left: ECIII. WT (*n* = 32): 4.13 ± 0.64 mV vs. *Gnb1*^I80T/+^ (*n* = 33): 3.30 ± 0.38 mV (*U* = 505, *p* = 0.7678). Center: CA3 Apical. WT (*n* = 31): 2.26 ± 0.33 mV vs. *Gnb1*^I80T/+^ (*n* = 32): 2.03 ± 0.36 mV (*U* = 406, *p* = 0.2185). Right: CA3 Basal. WT (*n* = 12): 3.61 ± 0.74 mV vs. *Gnb1*^I80T/+^ (*n* = 10): 1.45 ± 0.47 mV (*U* = 24, *p* = 0.0192). ***D***, Slow GABA_B_ IPSP area. Left: ECIII. WT (*n* = 33): 0.21 ± 0.04 mV * s vs. *Gnb1*^I80T/+^ (*n* = 34): 0.11 ± 0.01 mV * s (*U* = 306, *p* = 0.0014). Center: CA3 Apical. WT (*n* = 32): 0.05 ± 0.01 mV * s vs. *Gnb1*^I80T/+^ (*n* = 33): 0.05 ± 0.01 mV * s (*U* = 481, *p* = 0.05418). Right: CA3 Basal. WT (*n* = 12): 0.09 ± 0.02 mV * s vs. *Gnb1*^I80T/+^ (*n* = 10): 0.07 ± 0.02 mV * s (*U* = 51, *p* = 0.5752). All data represented as mean ± SEM. Statistics reflect Mann-Whitney U-tests. Asterisks indicate *p* < 0.05.

We next investigated the temporal summation of excitatory and inhibitory synaptic inputs by stimulating afferents with multiple pulses at varying frequencies. During repetitive action potential firing, synaptic input summation can either facilitate or depress in a frequency-dependent manner due to both short-term adaptation of presynaptic neurotransmitter release (Markram et al., 1998; Pouille and Scanziani, 2004; Klyachko and Stevens, 2006) and dynamics of postsynaptic voltage-dependent ion channels, including NMDA-type glutamate receptors, A-type potassium channels and hyperpolarization-activated cyclic nucleotide-gated (HCN) channels (Johnston et al., 1996; Magee, 2000; Losonczy and Magee, 2006). We observed pathway-specific changes in the frequency-dependent recruitment of synaptic excitation and inhibition in *Gnb1*^I80T/+^ neurons (Fig. 5). At the ECIII pathway to distal apical dendrites, both excitation and fast inhibition were enhanced at high stimulus frequencies (low ISIs), while at the CA3 pathway to basal dendrites, excitation, fast inhibition and slow inhibition were all decreased at high frequencies (Fig. 5). Quantitatively, the changes observed at basal dendritic synapses resulted in a larger ratio of excitation to inhibition (“E/I Imbalance,” Supp. Fig. S4). When compared to the expected amplitude of compound EPSPs predicted by linear summation of measured single-pulse EPSPs (Losonczy and Magee, 2006; Milstein et al., 2015) (Fig. 6A, see Methods), measured compound EPSPs were larger in amplitude across all stimulation pathways in both genotypes (Fig. 6A,B), consistent with engagement of pre- and post-synaptic facilitation mechanisms. However, this degree of supralinear summation was specifically increased in *Gnb1*^I80T/+^ neurons at the ECIII pathway (Fig. 6A,B). Taken together, these data suggest that decreased synaptic inhibition (Fig. 4C,D and 5C,D) and increased excitatory input summation (Fig. 6B) in *Gnb1*^I80T/+^ neurons may contribute to hyperexcitability of neuronal circuits in GNB1-E.

**Figure 5.**
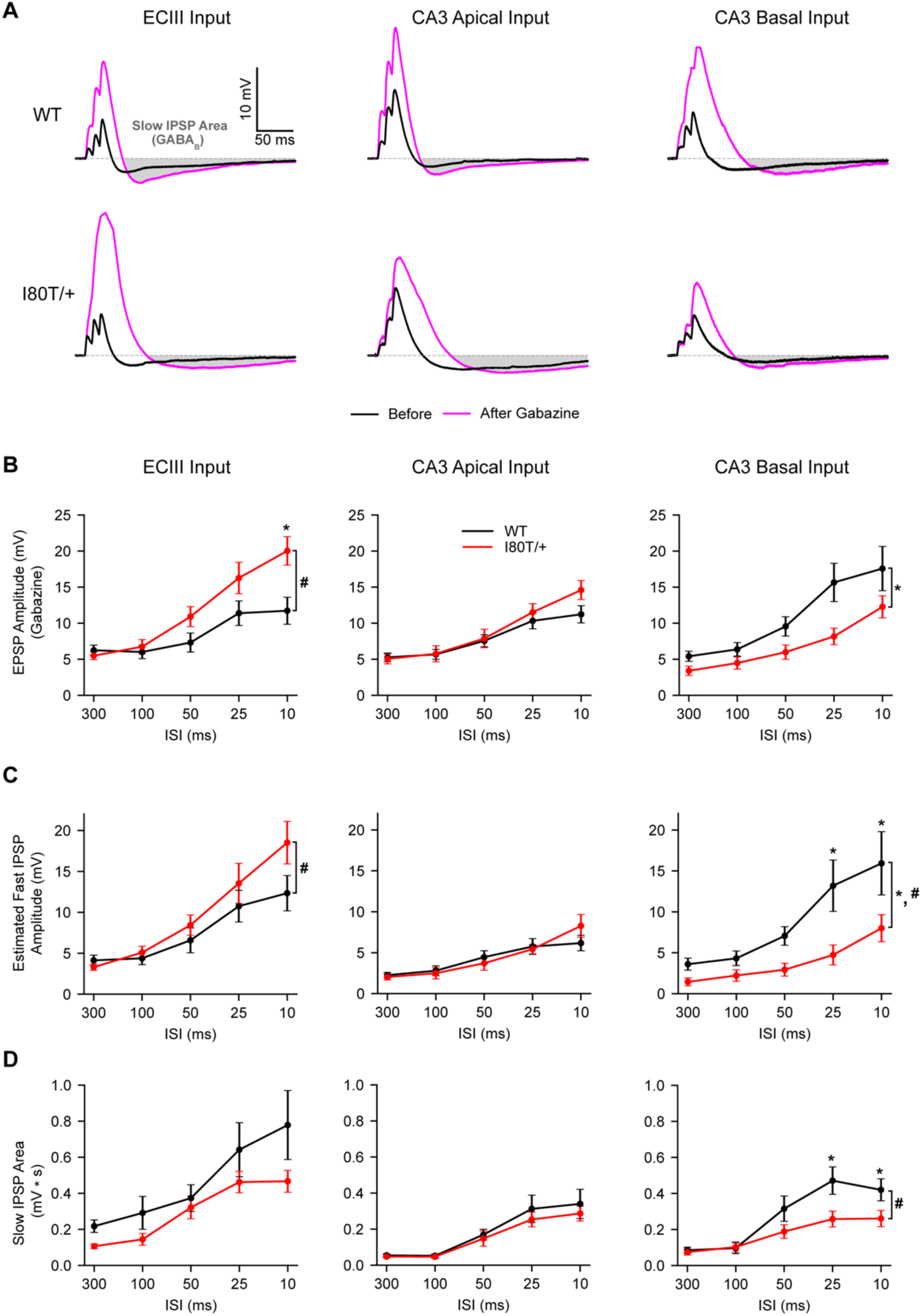
***A***, Example compound post-synaptic potentials (PSP) recorded before (black) and after application of gabazine (pink) during stimulation with a 10 ms ISI, allowing estimation of excitatory (EPSP), fast inhibitory (IPSP), and slow IPSP components (grey shaded area). Shown are representative examples from WT (top) and *Gnb1*^I80T/+^ (bottom) for stimulation of each of three input pathways. ***B***, EPSP amplitude. Left: ECIII (WT: *n* = 12–15, *Gnb1*^I80T/+^: *n* = 11–13). Type III ANOVA; main effect of genotype: *F*(1,21.0) *=* 3.689*, p =* 0.0684; genotype × ISI interaction effect: *F*(4,84.0) = 6.177, *p* = 0.0002; post-hoc t-tests: 10 ms ISI: *t*(39.3) = 3.791, *p* = 0.0025; other ISIs: *t*(39.3) = 0.043–2.220, *p* = 0.081–0.966 after FDR correction. Center: CA3 Apical (WT: *n* = 12–14, *Gnb1*^I80T/+^: *n* = 10–12). Type III ANOVA; main effect of genotype: *F*(1,20.0) = 1.004, *p* = 0.3284; genotype × ISI interaction effect: *F*(4,80.0) = 2.079, *p* = 0.0912. Right: CA3 Basal (WT: *n* = 11–12, *Gnb1*^I80T/+^: *n* = 10). Type III ANOVA; main effect of genotype: *F*(1,19.0) = 4.711, *p* = 0.0429; genotype × ISI interaction effect: *F*(4,76.0) = 2.476, *p* = 0.0512. ***C***, Estimated fast GABA_A_ IPSP amplitude (same neurons as ***B***). Left: ECIII. Type III ANOVA; main effect of genotype: *F*(1,19.0) = 0.4146, *p* = 0.5274; genotype × ISI interaction effect: *F*(4,76.0) = 2.875, *p* = 0.0284; post-hoc t-tests: all ISIs: *t*(34.5) = 0.023–2.058, *p* = 0.236–0.982 after FDR correction. Center: CA3 Apical. Type III ANOVA; main effect of genotype: *F*(1,19.0) = 0.1075, *p* = 0.7466; genotype × ISI interaction effect: *F*(4,76.0) = 2.413, *p* = 0.0562. Right: CA3 Basal. Type III ANOVA; main effect of genotype: *F*(1,19.0) = 5.445, *p* = 0.0308; genotype × ISI interaction effect: *F*(4,76.0) = 2.960, *p* = 0.0250; post-hoc t-tests: ISI 10 ms: *t*(43.4) = 3.126, *p* = 0.0097; ISI 25 ms: *t*(43.4) = 3.052, *p* = 0.0097; other ISIs: *t*(43.4) = 0.731–1.525, *p* = 0.224–0.469 after FDR correction. ***D***, Slow GABA_B_ IPSP area. Left: ECIII. Type III ANOVA; main effect of genotype: *F*(1,21.0) = 1.617, *p* = 0.2174; genotype × ISI interaction effect: *F*(4,84.0) = 1.625, *p* = 0.1755. Center: CA3 Apical. Type III ANOVA; main effect of genotype: *F*(1,20.0) = 0.1883, *p* = 0.6690; genotype × ISI interaction effect: *F*(4,80.0) = 0.4927, *p* = 0.7411. Right: CA3 Basal. Type III ANOVA; main effect of genotype: *F*(1,19.0) = 3.333, *p* = 0.0837; genotype × ISI interaction effect: *F*(4,76.0) = 5.011, *p* = 0.0012; post-hoc t-tests: ISI 10 ms: *t*(37.9) = 2.524, *p* = 0.0398; ISI 25 ms: *t*(37.9) = 3.029, *p* = 0.0220; other ISIs: *t*(37.9) = 0.013–1.923, *p* = 0.103–0.990 after FDR correction. All data represented as mean ± SEM. Statistics were performed using linear mixed effects models (see Methods). Asterisks with brackets indicate main effects with *p* < 0.05. Hashtags with brackets indicate interaction effects with *p* < 0.05. Asterisks above individual ISIs indicate post-hoc comparisons with *p* < 0.05 after FDR correction.

**Figure 6.**
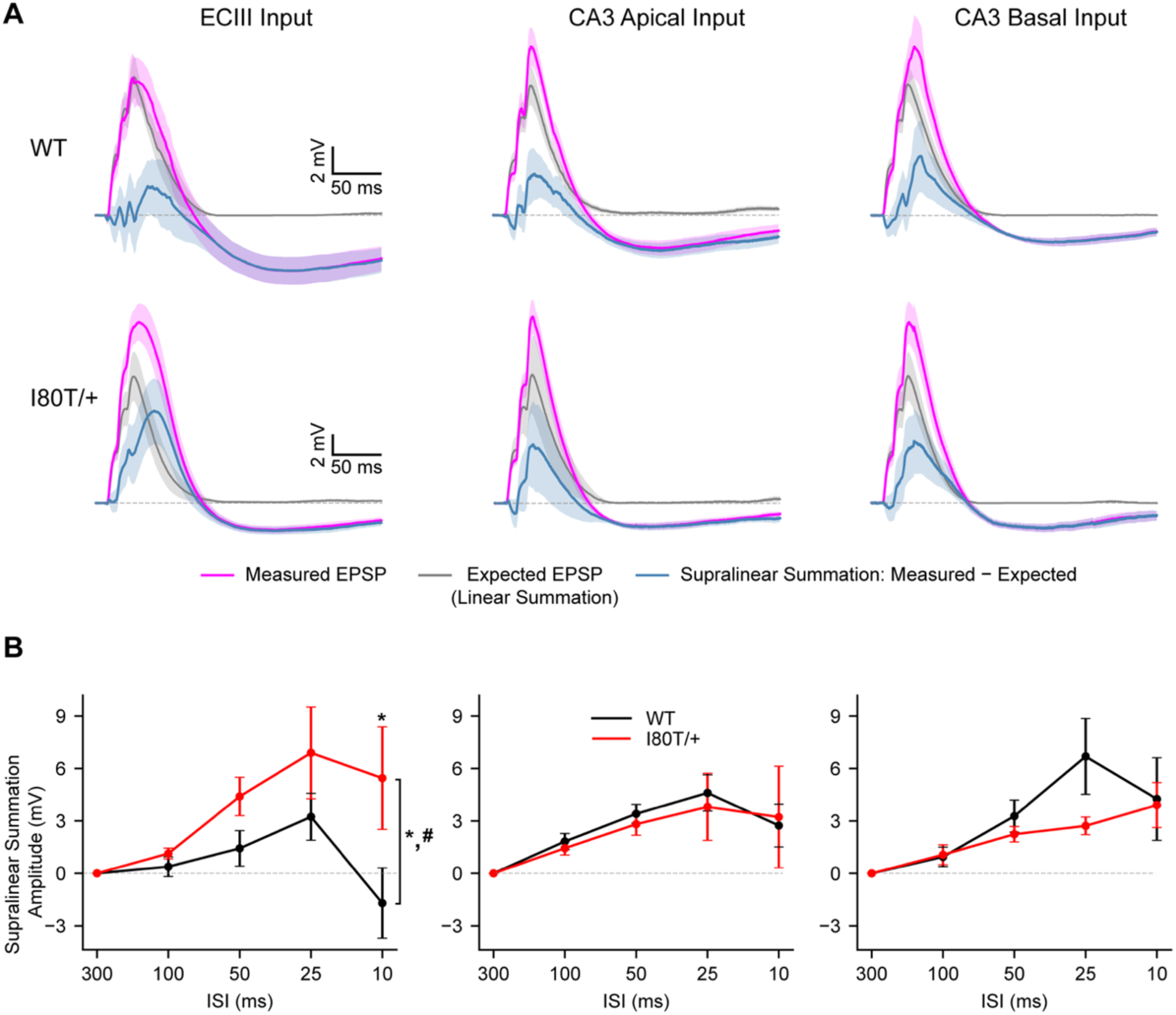
***A***, Traces include a compound EPSP measured during stimulation with a 10 ms ISI (pink), the compound EPSP expected from linear summation (grey, see Methods), and the difference trace (blue), reflecting the degree of supralinear summation. Shown are traces averaged across WT (top) or *Gnb1*^I80T/+^ (bottom) neurons for stimulation of each of three input pathways. Shading indicates SEM across neurons. ***B***, Supralinear summation is quantified. Left: ECIII (WT: *n* = 12–15, *Gnb1*^I80T/+^: *n* = 11–13 neurons). Type III ANOVA; main effect of genotype: *F*(1,76.9) = 9.852, *p* = 0.0024; genotype × ISI interaction effect: *F*(4,132.6) = 4.292, *p* = 0.0027; post-hoc t-tests: ISI 10 ms: *t*(156.2) = 4.158, *p* = 0.0003; other ISIs: *t*(131.4–155.7) = 0.000–2.201, *p* = 0.073–1.000 after FDR correction. Center: CA3 Apical (WT: *n* = 12– 14, *Gnb1*^I80T/+^: *n* = 10–12 neurons). Type III ANOVA; main effect of genotype: *F*(1,71.8) = 0.1489, *p* = 0.7007; genotype × ISI interaction effect: *F*(4,126.1) = 0.2038, *p* = 0.9359. Right: CA3 Basal (WT: *n* = 11–12, *Gnb1*^I80T/+^: *n* = 10 neurons). Type III ANOVA; main effect of genotype: *F*(1,19.8) = 0.7228, *p* = 0.4054; genotype × ISI interaction effect: *F*(4,78.9) = 1.196, *p* = 0.3192. All data represented as mean ± SEM. Statistics were performed using linear mixed effects models (see Methods). Asterisks with brackets indicate main effects with *p* < 0.05. Hashtags with brackets indicate interaction effects with *p* < 0.05. Asterisks above individual ISIs indicate post-hoc comparisons with *p* < 0.05 after FDR correction.

### Increased dendritic excitability in *Gnb1^I80T/+^* neurons

Having observed that supralinear summation of trains of synaptic inputs to distal dendrites is enhanced in *Gnb1*^I80T/+^ neurons, we next investigated whether this results in increased activation of voltage-gated ion channels in dendrites in response to physiologically relevant patterns of synaptic input. In hippocampal and cortical pyramidal neurons, coincidence of proximal and distal synaptic inputs can evoke a special type of regenerative depolarizing event called a dendritic calcium spike, or plateau potential, which reflects the activation of voltage-gated sodium, calcium and NMDA-type glutamate receptor channels (Takahashi and Magee, 2009; Larkum, 2013; Bittner et al., 2015; Milstein et al., 2015). In the rodent hippocampus, these events naturally occur during spatial foraging behavior when synchronous punctuated bursts of synaptic inputs are activated rhythmically in phase with a prominent 4–10 Hz theta oscillation (Buzsaki, 2002; Mizuseki et al., 2009; Buzsaki and Moser, 2013; Bittner et al., 2015). In *ex vivo* hippocampal slices, dendritic calcium spikes can be evoked with synchronous stimulation of distal ECIII and proximal CA3 inputs in a theta burst pattern (Takahashi and Magee, 2009; Milstein et al., 2015) (see Methods). When recording intracellularly in the cell soma, these events manifest as burst firing and large spike after-depolarizations that outlast the synaptic stimulus (Takahashi and Magee, 2009; Bittner et al., 2015; Milstein et al., 2015). Here we found that *Gnb1*^I80T/+^ neurons respond to theta burst stimulation of synaptic inputs by evoking longer duration dendritic calcium spikes compared to WT neurons (Fig. 7A–C). In fact, while WT neurons required both distal ECIII and proximal CA3 inputs to be coincidently stimulated to evoke dendritic calcium spikes, *Gnb1*^I80T/+^ neurons often produced dendritic calcium spikes in response to stimulation of just the ECIII pathway alone (Fig. 7A–C). Consistent with regenerative amplification by voltage-gated cation currents, supralinear input summation was increased in *Gnb1*^I80T/+^ neurons in response to theta burst stimulation of either distal ECIII or proximal CA3 pathways alone in addition to synchronous stimulation of both pathways (Fig. 7D,E). This observed increase in dendritic excitability is expected to increase somatic burst firing in response to dendritic synaptic input, which could contribute to increased seizure susceptibility in GNB1-E.

**Figure 7.**
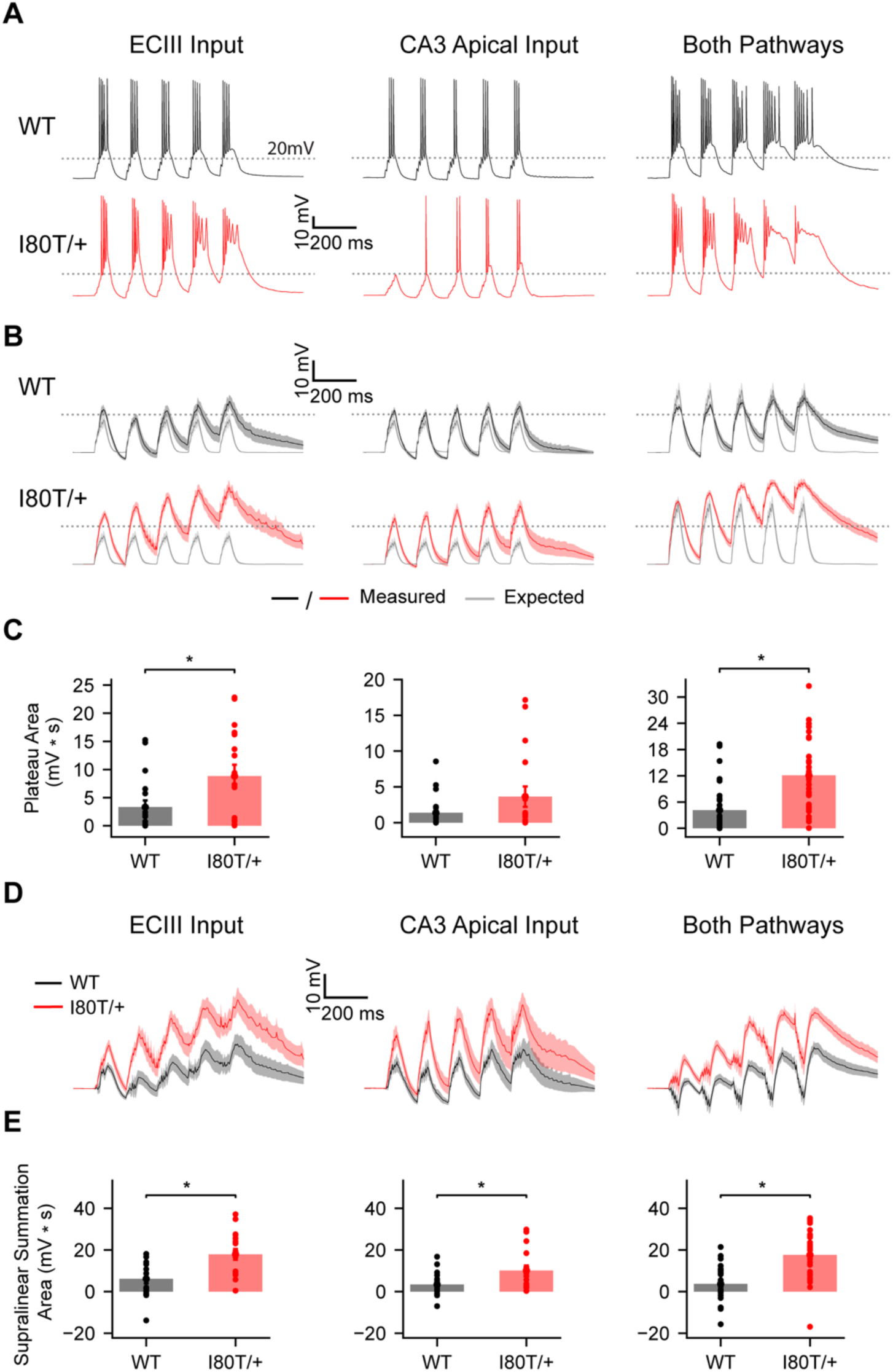
***A***, Example traces show compound EPSPs in response to theta-burst stimulation (see Methods). Shown are representative examples from WT (top) and *Gnb1*^I80T/+^ (bottom) neurons for stimulation of either ECIII inputs alone (left), CA3 inputs alone (center), or both pathways simultaneously (right). ***B***, Traces from experiments in ***A*** were processed by offline removal of fast sodium spikes, and averaged across WT (top, black) or *Gnb1*^I80T/+^ (bottom, red) neurons. Compound EPSPs expected from linear summation are shown in grey (see Methods). Shading indicates SEM across neurons. ***C***, Plateau area is quantified as the area of traces in ***B*** above a threshold of 20 mV above resting membrane potential (grey dotted line in ***B*** and ***C***). Left: ECIII. WT (*n* = 18): 3.30 ± 1.20 mV * s vs. *Gnb1*^I80T/+^ (*n* = 17): 8.83 ± 2.01 mV * s (*U* = 81, *p* = 0.0181). Center: CA3 Apical. WT (*n* = 19): 1.38 ± 0.54 mV * s vs. *Gnb1*^I80T/+^ (*n* = 17): 3.64 ± 1.42 mV * s (*U* = 122, *p* = 0.2130). Right: Both pathways. WT (*n* = 29): 4.15 ± 1.06 mV * s vs. *Gnb1*^I80T/+^ (*n* = 31): 12.09 ± 1.45 mV * s (*U* = 161, *p* < 0.0001). ***D***, Supralinear summation traces were generated using the same procedure as Fig. 6A. Shown are difference traces (Measured - Expected) averaged across WT (black) or *Gnb1*^I80T/+^ (red) neurons for stimulation of either ECIII inputs alone (left), CA3 inputs alone (center), or both pathways simultaneously (right). Shading indicates SEM across neurons. ***E***, Supralinear summation is quantified as the signed net area above or below zero in the traces in ***D***. Left: ECIII. WT (*n* = 18): 6.13 ± 2.13 mV * s vs. *Gnb1*^I80T/+^ (*n* = 17): 17.91 ± 2.59 mV * s (*U* = 61, *p* = 0.0025). Center: CA3 Apical. WT (*n* = 19): 3.39 ± 1.28 mV * s vs. *Gnb1*^I80T/+^ (*n* = 17): 10.15 ± 2.32 mV * s (*U* = 83, *p* = 0.0134). Right: Both pathways. WT (*n* = 29): 3.70 ± 1.52 mV * s vs. *Gnb1*^I80T/+^ (*n* = 31): 17.58 ± 2.14 mV * s (*U* = 145, *p* < 0.0001). All data represented as mean ± SEM. Statistics reflect Mann-Whitney U-tests. Asterisks indicate *p* < 0.05.

### Rescue of dendritic hyperexcitability in *Gnb1*^I80T/+^ neurons with a GIRK agonist

Taken together, the above-mentioned dendritic hyperexcitability (Fig. 7) and decrease in slow IPSP area (Fig. 4D and 5D) implicates disrupted activation of inhibitory GIRK channels by synaptically released GABA onto GABA_B_ receptors in the dendrites of pyramidal neurons (Fig. 8A). We next sought to directly test these mechanisms pharmacologically. First, we applied baclofen, a selective agonist of GABA_B_ receptors (Misgeld et al., 1995), and observed a large (∼6 mV) hyperpolarization of resting membrane potential in WT neurons that was substantially reduced (∼2 mV) in *Gnb1*^I80T/+^ neurons (Fig. 8B,C), corroborating a specific deficit in signaling downstream of GABA_B_ receptors. Next, we applied ML297, a selective activator of GIRK channels (Kaufmann et al., 2013; Wydeven et al., 2014; Huang et al., 2018), and measured slow IPSP area in response to synaptic stimulation (see Methods). This procedure is expected to activate GIRK channels and occlude any additional transient activation by GABA_B_ receptors (Dutar et al., 2000). We observed a reduction in slow IPSP area in both WT and *Gnb1*^I80T/+^ neurons (Fig. 8D,E), suggesting that the remaining slow GABA_B_ receptor IPSP in heterozygous *Gnb1*^I80T/+^ mice is still mediated by GIRK channels, and that the function of the non-mutated allele of *Gnb1* is still at least partially intact.

**Figure 8.**
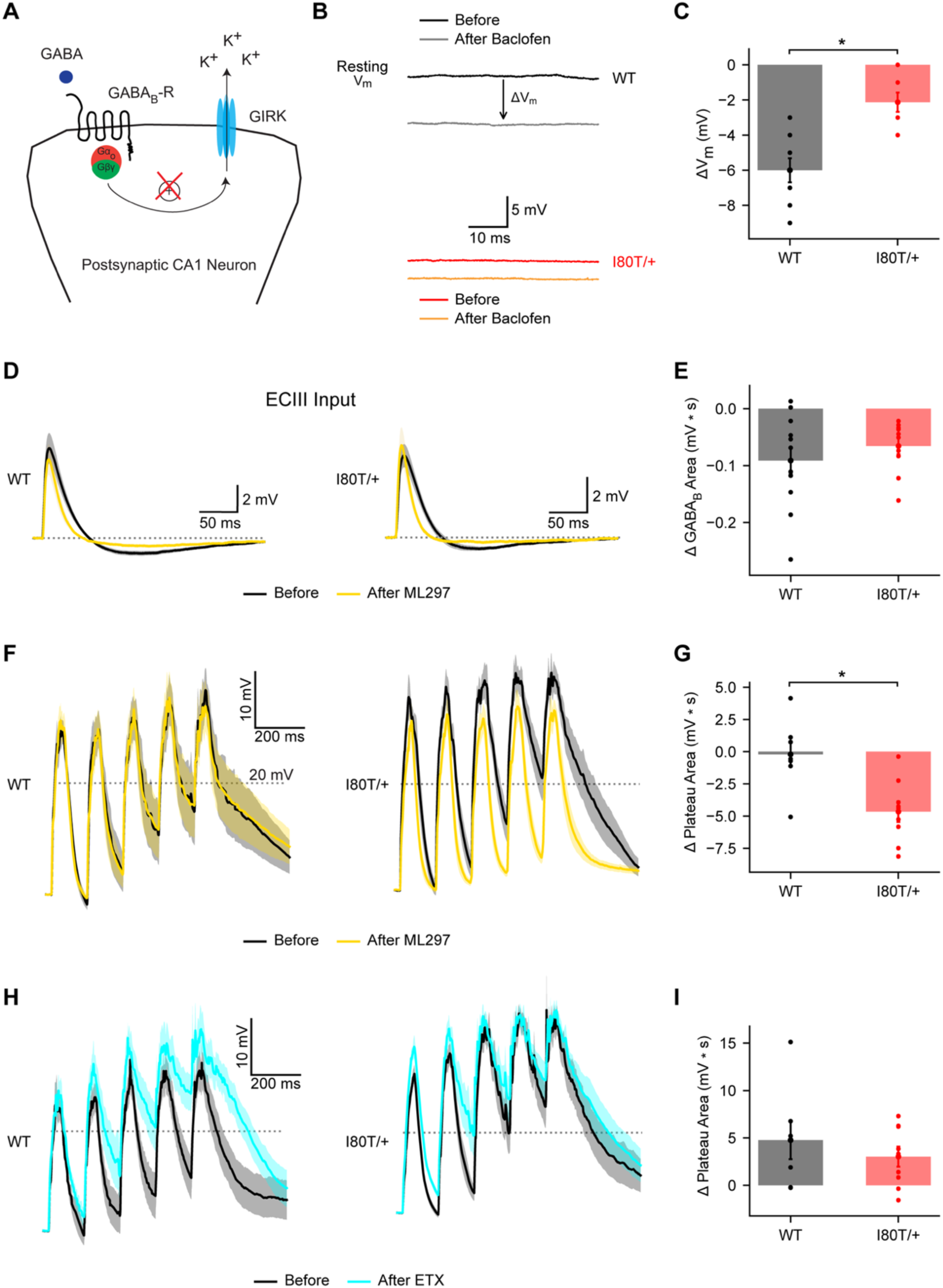
***A***, Diagram illustrates GABA_B_ receptors activating GIRK channels via a G protein signaling cascade that includes Gβ_1_ proteins. ***B***, Example traces show resting membrane potential recorded before and after application of the GABA_B_ agonist baclofen. Shown are recordings from WT (top, before: black, after: grey) and Gnb1^I80T/+^ (bottom, before: red, after: yellow) neurons. ***C***, Changes in membrane potential from experiments in ***B*** are quantified. WT (*n* = 9): −6.00 ± 0.69 mV vs. Gnb1^I80T/+^ (*n* = 8): −2.13 ± 0.55 mV (*U* = 68, *p* = 0.0020). ***D***, Traces show responses to stimulation of ECIII inputs before (black) and after (yellow) application of the GIRK activator ML297. Traces are averaged across 3 pulses stimulated with a 300 ms ISI, and then averaged across neurons (WT: left, *Gnb1*^I80T/+^: right). ***E***, Changes in slow GABA_B_ IPSP area from experiments in ***D*** are quantified. WT (*n* = 12): 0.091 ± 0.023 mV * s vs. Gnb1^I80T/+^ (*n* = 10): 0.066 ± 0.014 mV * s (*U* = 70, *p* = 0.5310). ***F***, Traces show responses to theta-burst stimulation of both ECIII and CA3 input pathways following offline removal of fast sodium spikes and averaging across neurons, similar to Fig. 6B. Shown are averaged traces from WT (left) or *Gnb1*^I80T/+^ (right) neurons before (black) and after (yellow) application of ML297. ***G***, Changes in plateau area from experiments in ***F*** are quantified. WT (*n* = 8): −0.21 ± 0.91 mV * s vs. *Gnb1*^I80T/+^ (*n* = 9): −4.66 ± 0.81 mV * s (*U* = 64, *p* = 0.0055). ***H***–***I***. Same as ***F***–***G*** but for application of the GIRK antagonist ethosuximide (before: black, after: cyan). WT (n=7): 4.76 ± 2.01 mV * s vs. *Gnb1*^I80T/+^ (*n* = 9): 3.03 ± 1.05 mV * s (*U* = 37, *p* = 0.6065). All data represented as mean ± SEM. Statistics reflect Mann-Whitney U-tests. Asterisks indicate *p* < 0.05.

We also measured dendritic calcium spiking before and after application of ML297 (Fig. 8F). While this had little effect in WT neurons where GIRK activation is likely already saturated by intact Gβ_1_ signaling, in *Gnb1*^I80T/+^ neurons this substantially reduced dendritic plateau area, rescuing the pathological hyperexcitability (Fig. 8F,G). The converse manipulation, application of the GIRK channel blocker ethosuximide (Kobayashi et al., 2009; Shalomov et al., 2025), resulted in increased dendritic plateau area in both WT and *Gnb1*^I80T/+^ neurons (Fig. 8H,I). This again suggests that some partial activation of GIRK channels by GABA_B_ receptors remains intact in *Gnb1*^I80T/+^ neurons, although it is also possible that ethosuximide increased dendritic excitability through off-target effects on calcium-activated potassium channels (Crunelli and Leresche, 2002). These results demonstrate that functional GIRK channels remain expressed at high levels in *Gnb1*^I80T/+^ neurons and suggest that their pharmacological activation has therapeutic potential to reduce neuronal hyperexcitability in GNB1-E caused by the p.I80T mutation.

## Discussion

In this study, we investigated the cellular, circuit, and synaptic mechanisms of neuronal dysfunction in a mouse model of GNB1 encephalopathy (GNB1-E) induced by p.I80T mutation of the *Gnb1* gene encoding the G protein subunit Gβ_1_. We found that heterozygous *Gnb1*^I80T/+^ mutation causes a deficit in synaptic inhibition and a pathological increase in dendritic excitability in mouse hippocampal neurons by disrupting activation of GIRK channels by synaptic GABA_B_ receptors. The GIRK activator drug ML297 suppressed this dendritic hyperexcitability and restored normal responses to synaptic afferent stimulation, providing preclinical evidence that targeting dendritic excitability could be a viable therapeutic strategy for GNB1-E (Xu et al., 2020; Zhao et al., 2020; Nguyen et al., 2024). Interestingly, while p.I80T is associated with epilepsy in humans and mice and causes GIRK loss of function, another common variant, p.K78R, is also associated with epilepsy, but causes GIRK gain of function (Petrovski et al., 2016; Hemati et al., 2018; Reddy et al., 2021; Colombo et al., 2023; Reddy et al., 2026). While the drug ethosuximide is used clinically for absence seizures due to its antagonism of calcium channels (Coulter et al., 1989), it also blocks GIRK channels and was shown to reduce seizures in *Gnb1*^K78R/+^ mice (Colombo et al., 2023; Shalomov et al., 2025). However, our results suggest that ethosuximide would in fact exacerbate increased dendritic excitability in the case of p.I80T, underscoring the necessity of patient-specific mechanism-based medicine.

While deficits in somatic excitability, E/I imbalance, and neuronal morphology are commonly observed in mouse models of epilepsy and neurodevelopmental disorders (George, 2004; Copf, 2016; Lee et al., 2017; Oyrer et al., 2018; Shao et al., 2019; Prem et al., 2020; Meisler et al., 2021; Prem et al., 2024; Xie et al., 2025), increasing evidence suggests that these disorders can also be associated with pathological changes to the unique computational properties of compartmentalized dendrites (Dyhrfjeld-Johnsen et al., 2008; Poolos and Johnston, 2012; Zhang et al., 2014; Spratt et al., 2019; Nelson and Bender, 2021; Brandalise et al., 2023). Interestingly, we observed that somatic excitability was counterintuitively decreased in *Gnb1*^I80T/+^ mice, possibly as a homeostatic compensation for the increase in dendritic excitability. The specific loss of dendritic inhibition and resulting increase in supralinear input summation at the long-range cortical inputs to the distal apical tuft dendrites of pyramidal neurons may have consequences for information coding and learning, as dendritic calcium spikes and their regulation by dendrite-targeting inhibitory interneurons have recently been identified as key mechanisms regulating long-term memory formation (Takahashi and Magee, 2009; Lovett-Barron et al., 2012; Xu et al., 2012; Larkum, 2013; Bittner et al., 2015; Milstein et al., 2015; Bittner et al., 2017; Grienberger et al., 2017; Magee and Grienberger, 2020; Grienberger and Magee, 2022; Rolotti et al., 2022; Li et al., 2024; Madar et al., 2025; Udakis et al., 2025; Xiao et al., 2025; Yaeger et al., 2025; Campbell et al., 2026; Magee, 2026; Vaasjo et al., 2026). We speculate that increased sensitivity to top-down instructive cortical inputs and increased plateau potential frequency and duration could cause plasticity to be induced too frequently and at inappropriate times, resulting in memory instability.

While here we focused on the hippocampus for its known roles in cognition and seizure susceptibility (Lenck-Santini and Scott, 2015), the absence epilepsy and motor deficits associated with GNB1-E could also result from additional disruptions in neuronal function in the thalamus, cortex, striatum, and/or cerebellum (Bosch-Bouju et al., 2013; Crunelli et al., 2020; Elder et al., 2025), meriting future investigation. Similar neurodevelopmental disorders and epilepsy also result from pathogenic variation in genes encoding other components of the inhibitory G protein signaling pathway, including *GNAI1* and *GNAO1* (Feng et al., 2017; Muir et al., 2021), highlighting the importance of understanding their roles in brain development and function, and continuing to develop mechanism-based treatments (Yu et al., 2019).

## Acknowledgments

This study is dedicated to K.R.M. We are grateful for resources and support provided by Jennifer Mulle and members of the Mulle Lab, and for funding provided by Rutgers Biomedical and Health Sciences (S.G., M.S.S., A.D.M.) and NIMH grants R01MH131296 (M.S.S.) and RF1MH135576 (S.G., A.V., A.R.G., A.D.M.).

## Supplementary Information

**Supplementary Figure 1.**
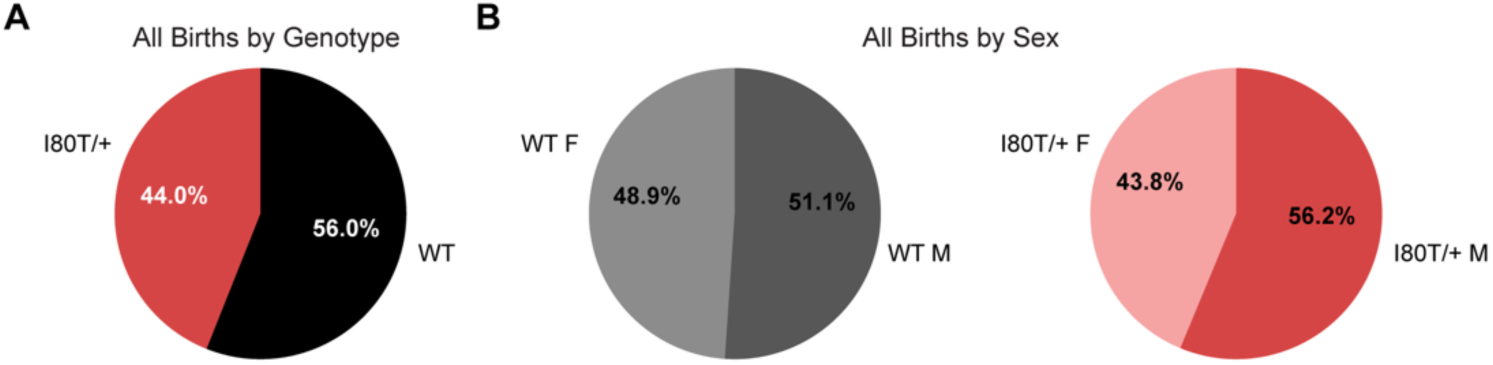
***A***, Pie chart shows the percentage of offspring of each genotype obtained from breeding heterozygous *Gnb1*^I80T/+^ and WT mice. ***B***, Pie charts show percentage of male and female offspring born of either WT (left) or *Gnb1*^I80T/+^ (right) genotype.

**Supplementary Figure 2.**
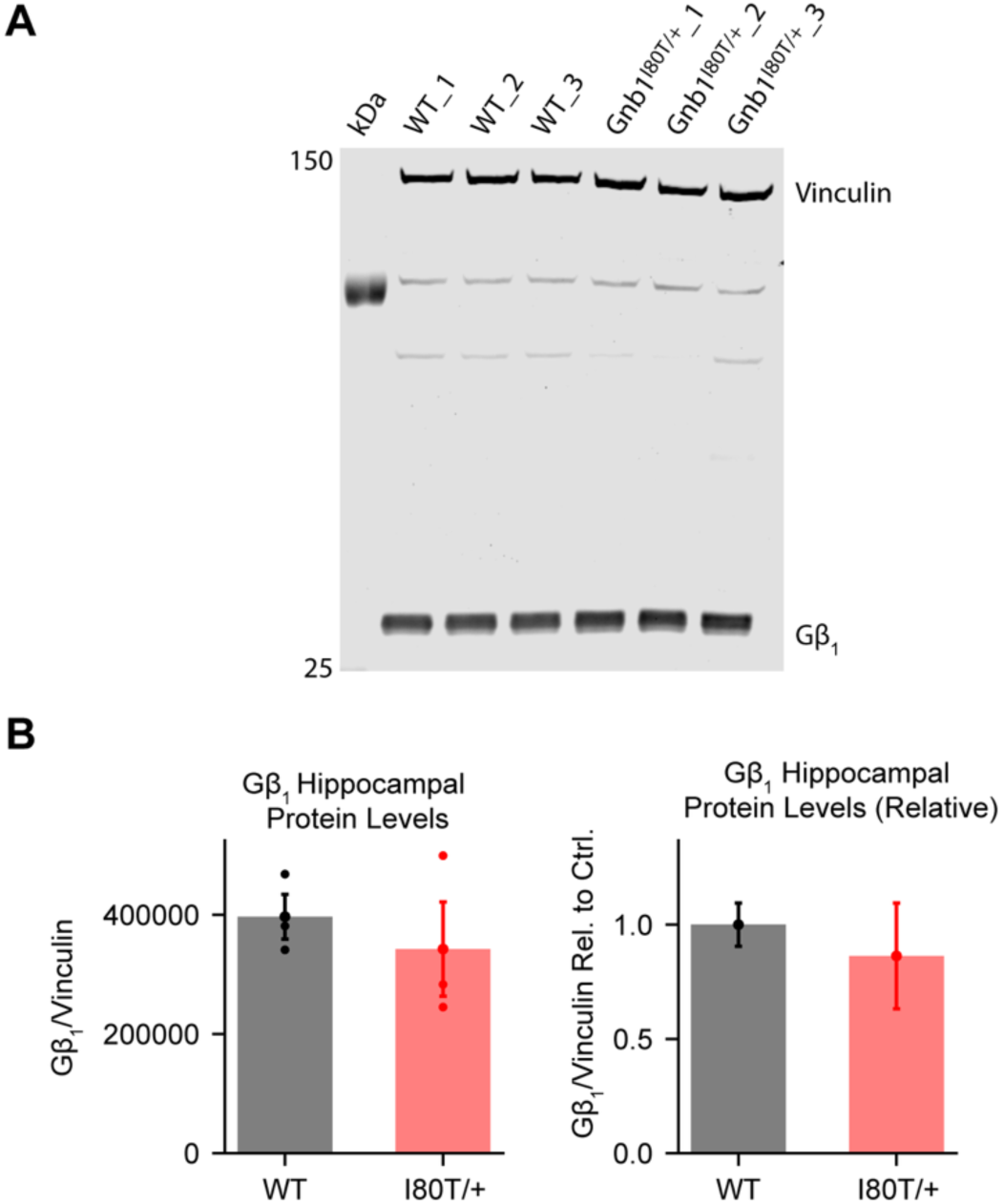
***A***, Western blot analysis of hippocampal protein lysates isolated from wild-type (WT) and *Gnb1*^I80T/+^ mice with antibodies against Gβ_1_ and Vinculin, used as a loading control (see Methods). ***B***, Left: Gβ_1_ protein levels from experiments in ***A*** are quantified as a ratio relative to Vinculin. WT (*n* = 3): 396666.67 ± 37489.26; *Gnb1*^I80T/+^ (*n* = 3): 342333.33 ± 79097.69 (*U* = 6, *p* = 0.7000). Right: Same as left, with protein ratios normalized to the average levels detected in WT samples. WT: 1.000 ± 0.095; *Gnb1*^I80T/+^: 0.863 ± 0.231 (*U* = 6, *p* = 0.7000). Data represented as mean ± SEM. Statistics reflect Mann-Whitney U-tests.

**Supplementary Figure 3.**
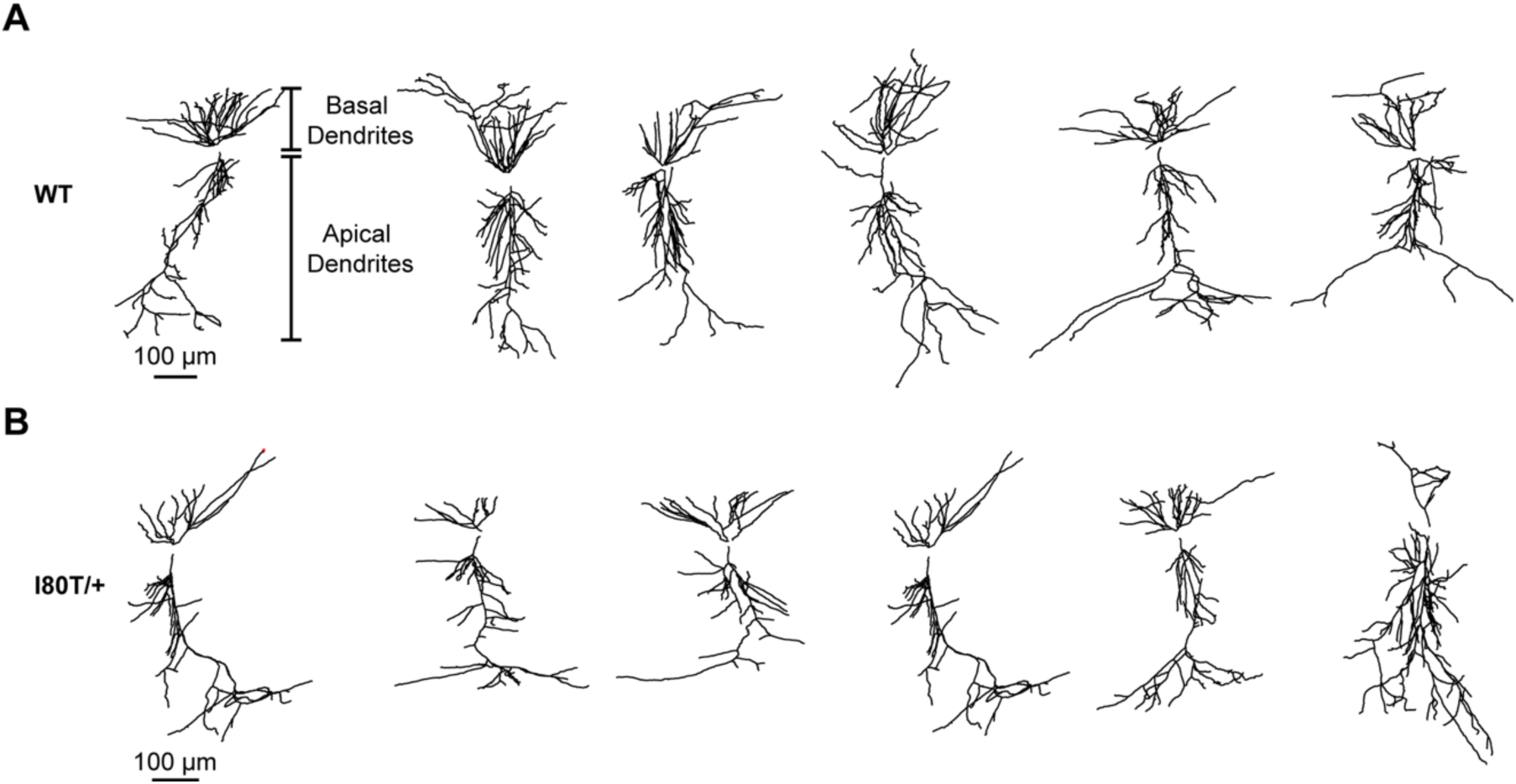
***A***, Representative histological reconstructions of wild-type (WT) neuron morphologies. ***B***, Same as ***A*** for *Gnb1*^I80T/+^ neurons.

**Supplementary Figure 4.**
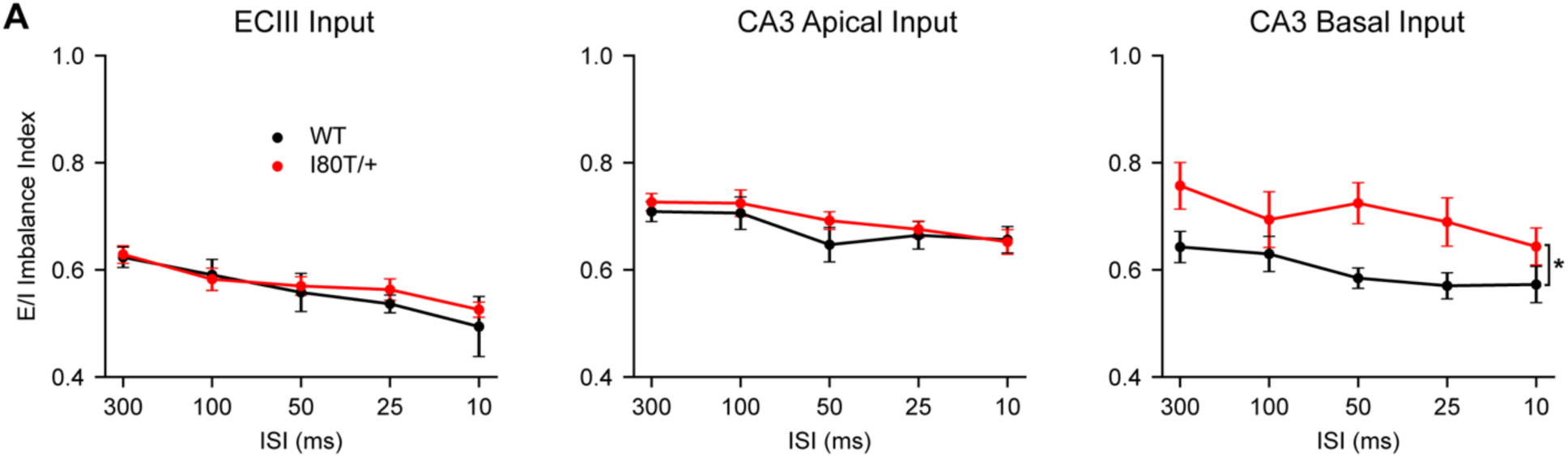
***A***, The ratio of excitation to inhibition is quantified as (EPSP/(EPSP + IPSP) (see Methods) from responses to stimulation of three input pathways (see Methods). Left: ECIII (WT: *n* = 12–15, *Gnb1*^I80T/+^: *n* = 11–13 neurons). Type III ANOVA; main effect of genotype: *F*(1, 73.5) *=* 0.047*, p* = 0.8290; genotype × ISI interaction effect: *F*(4, 105.1) = 0.116, *p* = 0.9767. Center: CA3 Apical (WT: *n* = 12–14, *Gnb1*^I80T/+^: *n* = 10–12). Type III ANOVA; main effect of genotype: *F*(1, 65.8) = 0.264, *p* = 0.6093; genotype × ISI interaction effect: *F*(4, 94.0) = 0.542, *p* = 0.7054. Right: CA3 Basal (WT: *n* = 11–12, *Gnb1*^I80T/+^: *n* = 10). Type III ANOVA; main effect of genotype: *F*(1, 20.1) = 6.551, *p* = 0.0187; genotype × ISI interaction effect: *F*(4, 79.1) = 0.911, *p* = 0.4620. All data represented as mean ± SEM. Statistics were performed using linear mixed effects models (see Methods). Asterisks with brackets indicate main effects with *p* < 0.05.

## References

Amaral DG, Witter MP (1989) The three-dimensional organization of the hippocampal formation: a review of anatomical data. Neuroscience 31:571–591.

Arshadi C, Gunther U, Eddison M, Harrington KIS, Ferreira TA (2021) SNT: a unifying toolbox for quantification of neuronal anatomy. Nat Methods 18:374–377.

Bates D, Mächler M, Bolker B, Walker S (2015) Fitting Linear Mixed-Effects Models Using lme4. Journal of Statistical Software 67:1–48.

Benjamini Y, Hochberg Y (1995) Controlling the false discovery rate: a practical and powerful approach to multiple testing. Journal of the Royal statistical society: series B (Methodological) 57:289–300.

Betke KM, Wells CA, Hamm HE (2012) GPCR mediated regulation of synaptic transmission. Prog Neurobiol 96:304–321.

Bittner KC, Milstein AD, Grienberger C, Romani S, Magee JC (2017) Behavioral time scale synaptic plasticity underlies CA1 place fields. Science 357:1033–1036.

Bittner KC, Grienberger C, Vaidya SP, Milstein AD, Macklin JJ, Suh J, Tonegawa S, Magee JC (2015) Conjunctive input processing drives feature selectivity in hippocampal CA1 neurons. Nat Neurosci 18:1133–1142.

Bloss EB, Cembrowski MS, Karsh B, Colonell J, Fetter RD, Spruston N (2016) Structured Dendritic Inhibition Supports Branch-Selective Integration in CA1 Pyramidal Cells. Neuron 89:1016–1030.

Booker SA, Vida I (2018) Morphological diversity and connectivity of hippocampal interneurons. Cell Tissue Res 373:619–641.

Bosch-Bouju C, Hyland BI, Parr-Brownlie LC (2013) Motor thalamus integration of cortical, cerebellar and basal ganglia information: implications for normal and parkinsonian conditions. Front Comput Neurosci 7:163.

Brager DH, Johnston D (2014) Channelopathies and dendritic dysfunction in fragile X syndrome. Brain Res Bull 103:11–17.

Brandalise F, Kalmbach BE, Cook EP, Brager DH (2023) Impaired dendritic spike generation in the Fragile X prefrontal cortex is due to loss of dendritic sodium channels. The Journal of Physiology 601:831–845.

Buzsaki G (2002) Theta oscillations in the hippocampus. Neuron 33:325–340.

Buzsaki G, Moser EI (2013) Memory, navigation and theta rhythm in the hippocampal-entorhinal system. Nat Neurosci 16:130–138.

Campbell EP, Martin L, Magee JC, Grienberger C (2026) Learning-dependent feedback by OLM interneurons shapes CA1 representations. bioRxiv.

Collins JM, Singh B, Zwick ME, Rosati G, Rigamonti M, Urdiales C, Eswaraka JR (2025) Using Machine Learning and Predictive Artificial Intelligence to Determine Cage Change Frequency for Mice Housed in Individually Ventilated Cages and Drive Vivarium Operational Efficiency. Journal of the American Association for Laboratory Animal Science 64:682–695.

Colombo S, Reddy HP, Petri S, Williams DJ, Shalomov B, Dhindsa RS, Gelfman S, Krizay D, Bera AK, Yang M (2023) Epilepsy in a mouse model of GNB1 encephalopathy arises from altered potassium (GIRK) channel signaling and is alleviated by a GIRK inhibitor. Frontiers in cellular neuroscience 17:1175895.

Copf T (2016) Impairments in dendrite morphogenesis as etiology for neurodevelopmental disorders and implications for therapeutic treatments. Neuroscience & Biobehavioral Reviews 68:946–978.

Coulter DA, Huguenard JR, Prince DA (1989) Characterization of ethosuximide reduction of low-threshold calcium current in thalamic neurons. Ann Neurol 25:582–593.

Crawley JN (1985) Exploratory behavior models of anxiety in mice. Neuroscience & Biobehavioral Reviews 9:37–44.

Crunelli V, Leresche L (2002) Block of thalamic T-type Ca2+ channels by ethosuximide is not the whole story. Epilepsy currents 2:53–56.

Crunelli V, Lőrincz ML, McCafferty C, Lambert RC, Leresche N, Di Giovanni G, David F (2020) Clinical and experimental insight into pathophysiology, comorbidity and therapy of absence seizures. Brain 143:2341–2368.

d’Isa R, Comi G, Leocani L (2021) Apparatus design and behavioural testing protocol for the evaluation of spatial working memory in mice through the spontaneous alternation T-maze. Scientific Reports 11:21177.

Da Silva JD, Costa MD, Almeida B, Lopes F, Maciel P, Teixeira-Castro A (2021) Case Report: A Novel GNB1 Mutation Causes Global Developmental Delay With Intellectual Disability and Behavioral Disorders. Front Neurol 12:735549.

Dascal N (2001) Ion-channel regulation by G proteins. Trends Endocrinol Metab 12:391–398.

Dascal N, Kahanovitch U (2015) The Roles of Gbetagamma and Galpha in Gating and Regulation of GIRK Channels. Int Rev Neurobiol 123:27–85.

DeFelipe J, Alonso-Nanclares L, Arellano JI (2002) Microstructure of the neocortex: comparative aspects. J Neurocytol 31:299–316.

Degro CE, Kulik A, Booker SA, Vida I (2015) Compartmental distribution of GABAB receptor-mediated currents along the somatodendritic axis of hippocampal principal cells. Front Synaptic Neurosci 7:6.

Dutar P, Nicoll RA (1988) A physiological role for GABAB receptors in the central nervous system. Nature 332:156–158.

Dutar P, Petrozzino JJ, Vu HM, Schmidt MF, Perkel DJ (2000) Slow synaptic inhibition mediated by metabotropic glutamate receptor activation of GIRK channels. J Neurophysiol 84:2284–2290.

Dyhrfjeld-Johnsen J, Morgan RJ, Foldy C, Soltesz I (2008) Upregulated H-current in hyperexcitable CA1 dendrites after febrile seizures. Front Cell Neurosci 2:2.

Elder C, Kerestes R, Opal P, Marchese M, Devinsky O (2025) The cerebellum in epilepsy. Epilepsia 66:1773–1792.

Endo W, Ikemoto S, Togashi N, Miyabayashi T, Nakajima E, Hamano S-i, Shibuya M, Sato R, Takezawa Y, Okubo Y, Inui T, Kato M, Sengoku T, Ogata K, Hamanaka K, Mizuguchi T, Miyatake S, Nakashima M, Matsumoto N, Haginoya K (2020) Phenotype–genotype correlations in patients with GNB1 gene variants, including the first three reported Japanese patients to exhibit spastic diplegia, dyskinetic quadriplegia, and infantile spasms. Brain and Development 42:199–204.

Fellous JM, Rudolph M, Destexhe A, Sejnowski TJ (2003) Synaptic background noise controls the input/output characteristics of single cells in an in vitro model of in vivo activity. Neuroscience 122:811–829.

Feng H, Sjogren B, Karaj B, Shaw V, Gezer A, Neubig RR (2017) Movement disorder in GNAO1 encephalopathy associated with gain-of-function mutations. Neurology 89:762–770.

Fino E, Packer AM, Yuste R (2013) The logic of inhibitory connectivity in the neocortex. Neuroscientist 19:228–237.

Ford CE, Skiba NP, Bae H, Daaka Y, Reuveny E, Shekter LR, Rosal R, Weng G, Yang CS, Iyengar R, Miller RJ, Jan LY, Lefkowitz RJ, Hamm HE (1998) Molecular basis for interactions of G protein betagamma subunits with effectors. Science 280:1271–1274.

Fox J, Weisberg S (2019) An R Companion to Applied Regression, Third Edition. Thousand Oaks CA: Sage.

Freund TF, Buzsaki G (1996) Interneurons of the hippocampus. Hippocampus 6:347–470.

George AL, Jr. (2004) Inherited Channelopathies Associated with Epilepsy. Epilepsy Curr 4:65–70.

Giorgi FM, Ceraolo C, Mercatelli D (2022) The R Language: An Engine for Bioinformatics and Data Science. Life 12:648.

Gouveia K, Hurst JL (2017) Optimising reliability of mouse performance in behavioural testing: the major role of non-aversive handling. Sci Rep 7:44999.

Grienberger C, Magee JC (2022) Entorhinal cortex directs learning-related changes in CA1 representations. Nature 611:554–562.

Grienberger C, Milstein AD, Bittner KC, Romani S, Magee JC (2017) Inhibitory suppression of heterogeneously tuned excitation enhances spatial coding in CA1 place cells. Nat Neurosci 20:417–426.

Gritz S, Milstein AD (2026) Code repository for analysis of GNB1 I80T hippocampal slice data. https://github.com/Milstein-Lab/GNB1_paper_analysis_and_figures.

Gulyas AI, Megias M, Emri Z, Freund TF (1999) Total number and ratio of excitatory and inhibitory synapses converging onto single interneurons of different types in the CA1 area of the rat hippocampus. J Neurosci 19:10082–10097.

Hemati P et al. (2018) Refining the phenotype associated with GNB1 mutations: Clinical data on 18 newly identified patients and review of the literature. Am J Med Genet A 176:2259–2275.

Huang Y, Zhang Y, Kong S, Zang K, Jiang S, Wan L, Chen L, Wang G, Jiang M, Wang X, Hu J, Wang Y (2018) GIRK1-mediated inwardly rectifying potassium current suppresses the epileptiform burst activities and the potential antiepileptic effect of ML297. Biomedicine & Pharmacotherapy 101:362–370.

Johnston D, Magee JC, Colbert CM, Cristie BR (1996) Active properties of neuronal dendrites. Annu Rev Neurosci 19:165–186.

Kaufmann K, Romaine I, Days E, Pascual C, Malik A, Yang L, Zou B, Du Y, Sliwoski G, Morrison RD, Denton J, Niswender CM, Daniels JS, Sulikowski GA, Xie XS, Lindsley CW, Weaver CD (2013) ML297 (VU0456810), the first potent and selective activator of the GIRK potassium channel, displays antiepileptic properties in mice. ACS Chem Neurosci 4:1278–1286.

Klyachko VA, Stevens CF (2006) Excitatory and feed-forward inhibitory hippocampal synapses work synergistically as an adaptive filter of natural spike trains. PLoS Biol 4:e207.

Kobayashi T, Hirai H, Iino M, Fuse I, Mitsumura K, Washiyama K, Kasai S, Ikeda K (2009) Inhibitory effects of the antiepileptic drug ethosuximide on G protein-activated inwardly rectifying K+ channels. Neuropharmacology 56:499–506.

Koyrakh L, Lujan R, Colon J, Karschin C, Kurachi Y, Karschin A, Wickman K (2005) Molecular and cellular diversity of neuronal G-protein-gated potassium channels. J Neurosci 25:11468–11478.

Kuznetsova A, Brockhoff PB, Christensen RHB (2017) lmerTest Package: Tests in Linear Mixed Effects Models. Journal of Statistical Software 82:1–26.

Lansdon LA, Saunders CJ (2021) Genotype-phenotype correlation in GNB1-related neurodevelopmental disorder: Potential association of p.Leu95Pro with cleft palate. Am J Med Genet A 185:1341–1343.

Larkum M (2013) A cellular mechanism for cortical associations: an organizing principle for the cerebral cortex. Trends Neurosci 36:141–151.

Lee E, Lee J, Kim E (2017) Excitation/Inhibition Imbalance in Animal Models of Autism Spectrum Disorders. Biol Psychiatry 81:838–847.

Lenck-Santini PP, Scott RC (2015) Mechanisms Responsible for Cognitive Impairment in Epilepsy. Cold Spring Harb Perspect Med 5.

Lenth RV, Piaskowski J (2026) emmeans: Estimated Marginal Means, aka Least-Squares Means. In.

Li Y, Briguglio JJ, Romani S, Magee JC (2024) Mechanisms of memory-supporting neuronal dynamics in hippocampal area CA3. Cell 187:6804–6819 e6821.

Lohmann K, Masuho I, Patil DN, Baumann H, Hebert E, Steinrucke S, Trujillano D, Skamangas NK, Dobricic V, Huning I, Gillessen-Kaesbach G, Westenberger A, Savic-Pavicevic D, Munchau A, Oprea G, Klein C, Rolfs A, Martemyanov KA (2017) Novel GNB1 mutations disrupt assembly and function of G protein heterotrimers and cause global developmental delay in humans. Hum Mol Genet 26:1078–1086.

Losonczy A, Magee JC (2006) Integrative properties of radial oblique dendrites in hippocampal CA1 pyramidal neurons. Neuron 50:291–307.

Lovett-Barron M, Turi GF, Kaifosh P, Lee PH, Bolze F, Sun XH, Nicoud JF, Zemelman BV, Sternson SM, Losonczy A (2012) Regulation of neuronal input transformations by tunable dendritic inhibition. Nat Neurosci 15:423–430, S421-423.

Luo L (2021) Architectures of neuronal circuits. Science 373:eabg7285.

Luscher C, Slesinger PA (2010) Emerging roles for G protein-gated inwardly rectifying potassium (GIRK) channels in health and disease. Nat Rev Neurosci 11:301–315.

Maccaferri G, Mangoni M, Lazzari A, DiFrancesco D (1993) Properties of the hyperpolarization-activated current in rat hippocampal CA1 pyramidal cells. J Neurophysiol 69:2129–2136.

Madar AD, Milstein AD, O’Dell TJ, Jain A, Clopath C, Sheffield MEJ (2025) Behavioral Timescale Synaptic Plasticity: A Burst in the Field of Learning and Memory. The Journal of Neuroscience 45:e1332252025.

Magee JC (2000) Dendritic integration of excitatory synaptic input. Nat Rev Neurosci 1:181–190.

Magee JC (2026) Behavioral timescale synaptic plasticity: properties, elements and functions. Nat Neurosci 29:520–534.

Magee JC, Grienberger C (2020) Synaptic Plasticity Forms and Functions. Annu Rev Neurosci 43:95–117.

Malik R, Johnston D (2017) Dendritic GIRK Channels Gate the Integration Window, Plateau Potentials, and Induction of Synaptic Plasticity in Dorsal But Not Ventral CA1 Neurons. J Neurosci 37:3940–3955.

Mark MD, Herlitze S (2000) G-protein mediated gating of inward-rectifier K+ channels. Eur J Biochem 267:5830–5836.

Markram H, Gupta A, Uziel A, Wang Y, Tsodyks M (1998) Information processing with frequency-dependent synaptic connections. Neurobiol Learn Mem 70:101–112.

Masala N, Pofahl M, Haubrich AN, Sameen Islam KU, Nikbakht N, Pasdarnavab M, Bohmbach K, Araki K, Kamali F, Henneberger C, Golcuk K, Ewell LA, Blaess S, Kelly T, Beck H (2023) Targeting aberrant dendritic integration to treat cognitive comorbidities of epilepsy. Brain 146:2399–2417.

Megias M, Emri Z, Freund TF, Gulyas AI (2001) Total number and distribution of inhibitory and excitatory synapses on hippocampal CA1 pyramidal cells. Neuroscience 102:527–540.

Meisler MH, Hill SF, Yu W (2021) Sodium channelopathies in neurodevelopmental disorders. Nat Rev Neurosci 22:152–166.

Milstein AD, Bloss EB, Apostolides PF, Vaidya SP, Dilly GA, Zemelman BV, Magee JC (2015) Inhibitory Gating of Input Comparison in the CA1 Microcircuit. Neuron 87:1274–1289.

Misgeld U, Bijak M, Jarolimek W (1995) A physiological role for GABAB receptors and the effects of baclofen in the mammalian central nervous system. Prog Neurobiol 46:423–462.

Mizuseki K, Sirota A, Pastalkova E, Buzsaki G (2009) Theta oscillations provide temporal windows for local circuit computation in the entorhinal-hippocampal loop. Neuron 64:267–280.

Muir AM et al. (2021) Variants in GNAI1 cause a syndrome associated with variable features including developmental delay, seizures, and hypotonia. Genet Med 23:881–887.

Nasvytis M, Čiauškaitė J, Jurkevičienė G (2024) GNB1 Encephalopathy: Clinical Case Report and Literature Review. Medicina 60:589.

Nelson AD, Bender KJ (2021) Dendritic Integration Dysfunction in Neurodevelopmental Disorders. Dev Neurosci 43:201–221.

Nguyen H, Glaaser IW, Slesinger PA (2024) Direct modulation of G protein-gated inwardly rectifying potassium (GIRK) channels. Front Physiol 15:1386645.

Okae H, Iwakura Y (2010) Neural tube defects and impaired neural progenitor cell proliferation in Gbeta1-deficient mice. Dev Dyn 239:1089–1101.

Oldham WM, Hamm HE (2008) Heterotrimeric G protein activation by G-protein-coupled receptors. Nat Rev Mol Cell Biol 9:60–71.

Oyrer J, Maljevic S, Scheffer IE, Berkovic SF, Petrou S, Reid CA (2018) Ion Channels in Genetic Epilepsy: From Genes and Mechanisms to Disease-Targeted Therapies. Pharmacol Rev 70:142–173.

Palmer L, Murayama M, Larkum M (2012) Inhibitory Regulation of Dendritic Activity in vivo. Front Neural Circuits 6:26.

Petreanu L, Mao T, Sternson SM, Svoboda K (2009) The subcellular organization of neocortical excitatory connections. Nature 457:1142–1145.

Petrovski S et al. (2016) Germline De Novo Mutations in GNB1 Cause Severe Neurodevelopmental Disability, Hypotonia, and Seizures. Am J Hum Genet 98:1001–1010.

Pinheiro J, Bates D, DebRoy S, Sarkar D, Team RC (2026) nlme: Linear and Nonlinear Mixed Effects Models. In.

Pires G, Leitner D, Drummond E, Kanshin E, Nayak S, Askenazi M, Faustin A, Friedman D, Debure L, Ueberheide B (2021) Proteomic differences in the hippocampus and cortex of epilepsy brain tissue. Brain Communications 3:fcab021.

Poolos NP, Johnston D (2012) Dendritic ion channelopathy in acquired epilepsy. Epilepsia 53 Suppl 9:32–40.

Pouille F, Scanziani M (2004) Routing of spike series by dynamic circuits in the hippocampus. Nature 429:717–723.

Prem S, Millonig JH, DiCicco-Bloom E (2020) Dysregulation of Neurite Outgrowth and Cell Migration in Autism and Other Neurodevelopmental Disorders. Adv Neurobiol 25:109–153.

Prem S, Dev B, Peng C, Mehta M, Alibutud R, Connacher RJ, St Thomas M, Zhou X, Matteson P, Xing J, Millonig JH, DiCicco-Bloom E (2024) Dysregulation of mTOR signaling mediates common neurite and migration defects in both idiopathic and 16p11.2 deletion autism neural precursor cells. eLife 13:e82809.

Qayum S, Petrin D, Tanny JC, Hebert TE (2026) Gbetagamma Signaling: Lessons Across the Cellular Multiverse. Annu Rev Pharmacol Toxicol 66:487–500.

Reddy HP, Yakubovich D, Keren-Raifman T, Tabak G, Tsemakhovich VA, Pedersen MH, Shalomov B, Colombo S, Goldstein DB, Javitch JA, Bera AK, Dascal N (2021) Encephalopathy-causing mutations in Gbeta(1) (GNB1) alter regulation of neuronal GIRK channels. iScience 24:103018.

Reddy HP, Ranjan V, Klo M, Shapiro G, Bassan H, Harel G, Heimer G, Ben Zeev B, Rabinski T, Vatine GD, Yaffe Y, Maoz BM, Bikovski L, Shomron N, Yakubovich DM, Rubinstein M, Dascal N (2026) Cross-species analysis of GNB1 I80T encephalopathy: conserved developmental, epileptic and neuronal transcriptome signatures. bioRxiv 2026.08.03.742477.

Reyes NGD, Di Luca DG, McNiven V, Lang AE (2023) Dystonia with myoclonus and vertical supranuclear gaze palsy associated with a rare GNB1 variant. Parkinsonism & Related Disorders 106:105239.

Rolotti SV, Ahmed MS, Szoboszlay M, Geiller T, Negrean A, Blockus H, Gonzalez KC, Sparks FT, Solis Canales AS, Tuttman AL, Peterka DS, Zemelman BV, Polleux F, Losonczy A (2022) Local feedback inhibition tightly controls rapid formation of hippocampal place fields. Neuron 110:783–794 e786.

Rothman JS, Silver RA (2018) NeuroMatic: An Integrated Open-Source Software Toolkit for Acquisition, Analysis and Simulation of Electrophysiological Data. Front Neuroinform 12:14.

Schindelin J, Arganda-Carreras I, Frise E, Kaynig V, Longair M, Pietzsch T, Preibisch S, Rueden C, Saalfeld S, Schmid B, Tinevez JY, White DJ, Hartenstein V, Eliceiri K, Tomancak P, Cardona A (2012) Fiji: an open-source platform for biological-image analysis. Nat Methods 9:676–682.

Schultz-Rogers L, Masuho I, Pinto e Vairo F, Schmitz CT, Schwab TL, Clark KJ, Gunderson L, Pichurin PN, Wierenga K, Martemyanov KA, Klee EW (2020) Haploinsufficiency as a disease mechanism in GNB1-associated neurodevelopmental disorder. Molecular Genetics & Genomic Medicine 8:e1477.

Shalomov B, Friesacher T, Yakubovich D, Combista JC, Reddy HP, Dabbah S, Bernsteiner H, Zangerl-Plessl EM, Stary-Weinzinger A, Dascal N (2025) Ethosuximide: Subunit- and Gbetagamma-dependent blocker and reporter of allosteric changes in GIRK channels. Br J Pharmacol 182:1704–1718.

Shao LR, Habela CW, Stafstrom CE (2019) Pediatric Epilepsy Mechanisms: Expanding the Paradigm of Excitation/Inhibition Imbalance. Children (Basel) 6.

Sholl DA (1953) Dendritic organization in the neurons of the visual and motor cortices of the cat. J Anat 87:387–406.

Signorini S, Liao YJ, Duncan SA, Jan LY, Stoffel M (1997) Normal cerebellar development but susceptibility to seizures in mice lacking G protein-coupled, inwardly rectifying K+ channel GIRK2. Proc Natl Acad Sci U S A 94:923–927.

Smirnov S, Paalasmaa P, Uusisaari M, Voipio J, Kaila K (1999) Pharmacological isolation of the synaptic and nonsynaptic components of the GABA-mediated biphasic response in rat CA1 hippocampal pyramidal cells. J Neurosci 19:9252–9260.

Spratt PWE, Ben-Shalom R, Keeshen CM, Burke KJ, Jr., Clarkson RL, Sanders SJ, Bender KJ (2019) The Autism-Associated Gene Scn2a Contributes to Dendritic Excitability and Synaptic Function in the Prefrontal Cortex. Neuron 103:673–685 e675.

Spruston N (2008) Pyramidal neurons: dendritic structure and synaptic integration. Nat Rev Neurosci 9:206–221.

Steinrucke S, Lohmann K, Domingo A, Rolfs A, Baumer T, Spiegler J, Hartmann C, Munchau A (2016) Novel GNB1 missense mutation in a patient with generalized dystonia, hypotonia, and intellectual disability. Neurol Genet 2:e106.

Szczałuba K, Biernacka A, Szymańska K, Gasperowicz P, Kosińska J, Rydzanicz M, Płoski R (2018) Novel GNB1 de novo mutation in a patient with neurodevelopmental disorder and cutaneous mastocytosis: Clinical report and literature review. European Journal of Medical Genetics 61:157–160.

Takahashi H, Magee JC (2009) Pathway interactions and synaptic plasticity in the dendritic tuft regions of CA1 pyramidal neurons. Neuron 62:102–111.

Tir S, Foster RG, Peirson SN (2025) Evaluation of the Digital Ventilated Cage(R) system for circadian phenotyping. Sci Rep 15:3674.

Tsuji M, Ikeda A, Tsuyusaki Y, Iai M, Kurosawa K, Kosaki K, Goto T (2023) Atypical clinical course in two patients with GNB1 variants who developed acute encephalopathy. Brain and Development 45:462–466.

Udakis M, Claydon MDB, Zhu HW, Oakes EC, Mellor JR (2025) Hippocampal OLM interneurons regulate CA1 place cell plasticity and remapping. Nat Commun 16:9912.

Vaasjo LO, Kotermanski SE, Patel T, Shi HJ, Machold R, Chamberland S (2026) Dendritic Inhibition Terminates Plateau Potentials in CA1 Pyramidal Neurons. J Neurosci 46.

Virtanen P et al. (2020) SciPy 1.0: fundamental algorithms for scientific computing in Python. Nat Methods 17:261–272.

Wydeven N, Marron Fernandez de Velasco E, Du Y, Benneyworth MA, Hearing MC, Fischer RA, Thomas MJ, Weaver CD, Wickman K (2014) Mechanisms underlying the activation of G-protein-gated inwardly rectifying K+ (GIRK) channels by the novel anxiolytic drug, ML297. Proc Natl Acad Sci U S A 111:10755–10760.

Xiao K, Li Y, Sullivan BJ, Li G, Magee JC (2025) Rapid neocortical network modifications via dendritic plateau potential induced plasticity. bioRxiv:2025.2011.2019.689338.

Xie K, Royer J, Rodriguez-Cruces R, Horwood L, Ngo A, Arafat T, Auer H, Sahlas E, Chen J, Zhou Y, Valk SL, Hong SJ, Frauscher B, Pana R, Bernasconi A, Bernasconi N, Concha L, Bernhardt BC (2025) Temporal Lobe Epilepsy Perturbs the Brain-Wide Excitation-Inhibition Balance: Associations with Microcircuit Organization, Clinical Parameters, and Cognitive Dysfunction. Adv Sci (Weinh) 12:e2406835.

Xu NL, Harnett MT, Williams SR, Huber D, O’Connor DH, Svoboda K, Magee JC (2012) Nonlinear dendritic integration of sensory and motor input during an active sensing task. Nature 492:247–251.

Xu Y, Cantwell L, Molosh AI, Plant LD, Gazgalis D, Fitz SD, Dustrude ET, Yang Y, Kawano T, Garai S, Noujaim SF, Shekhar A, Logothetis DE, Thakur GA (2020) The small molecule GAT1508 activates brain-specific GIRK1/2 channel heteromers and facilitates conditioned fear extinction in rodents. J Biol Chem 295:3614–3634.

Yaeger CE, Mojica Soto-Albors R, Liu W, Beltramini A, Harnett MT (2025) Plateau potentials are instructive signals for behavioral timescale synaptic plasticity in the neocortex. bioRxiv:2025.2011.2007.687250.

Yu Y, Nguyen DT, Jiang J (2019) G protein-coupled receptors in acquired epilepsy: Druggability and translatability. Prog Neurobiol 183:101682.

Zamponi GW, Currie KP (2013) Regulation of Ca(V)2 calcium channels by G protein coupled receptors. Biochim Biophys Acta 1828:1629–1643.

Zhang Y, Bonnan A, Bony G, Ferezou I, Pietropaolo S, Ginger M, Sans N, Rossier J, Oostra B, LeMasson G, Frick A (2014) Dendritic channelopathies contribute to neocortical and sensory hyperexcitability in Fmr1(-/y) mice. Nat Neurosci 17:1701–1709.

Zhao Y, Ung PM, Zahoranszky-Kohalmi G, Zakharov AV, Martinez NJ, Simeonov A, Glaaser IW, Rai G, Schlessinger A, Marugan JJ, Slesinger PA (2020) Identification of a G-Protein-Independent Activator of GIRK Channels. Cell Rep 31:107770.

